# *Toxoplasma gondii* Fatty Acid Elongases are Important for Virulence and Persistence

**DOI:** 10.64898/2026.07.29.741455

**Authors:** Daniel Walsh, Leandro Tana-Hernandez, Yoshiki Yamaryo-Botté, Saniya Crouch, Ving-Yun Li, Leandro Lemgruber, David Smith, Cyrille Y. Botté, Maria E. Francia, Lilach Sheiner

**Author notes:** Equal contribution (Cyrille Y. Botté, Maria E. Francia, Lilach Sheiner).

## Abstract

*Toxoplasma gondii* is an apicomplexan parasite which can infect a diverse range of warm-blooded host species. This promiscuous lifestyle requires metabolic flexibility yet our understanding of the metabolic pathways in this parasite remains incomplete. This gap includes the endoplasmic reticulum localised fatty acid elongation (FAE) pathway, which is responsible for synthesis of long (LCFA) and very long chain fatty acids (VLCFA), thus contributing to critical functions such as lipid storage and membrane biogenesis. Analyses of FAE in the commonly studied type I *T. gondii* strain revealed its contribution to the production of LCFA and VLCFA *in vitro*. However, the cystogenic type II *T. gondii,* which is responsible for most toxoplasmosis cases worldwide, was not studied. These type II parasites can readily differentiate between virulent and immune-evasive stages whereas type I cannot. This ability to readily transition between life stages, which are metabolically fundamentally different, prompts the question of whether the FAE pathway may play different roles in their distinct phenotype. Here we address this question, first confirming the role of the FAE enzymes, TgELO-A & B, in the production of LCFA and VLCFA in a type II *T. gondii* strain. Next, we demonstrated that their deletion leads to defects in replication and invasion when grown under lipid-restricted conditions. The mutants were further shown to be attenuated in mice as demonstrated by extended survival. Importantly, FAE enzyme deletion caused reduced brain cyst burdens. These findings highlight the importance of FAE for parasite survival under metabolic stress and *in vivo*.

**Importance:** *Toxoplasma gondii* infects a wide range of hosts and persists for life in multiple tissues, making metabolic flexibility essential for its success. Fatty acid elongation (FAE), mediated by elongase (ELO) enzymes, enables the parasite to modify both de novo synthesised and host-scavenged fatty acids to support membrane biogenesis, lipid storage, and adaptation to changing nutrient environments. Previous studies in the type I strain established the importance of FAE for parasite metabolism, but its role in the clinically relevant, cystogenic type II lineage—which causes most cases of human toxoplasmosis and forms long-lived, immune-evasive bradyzoite cysts— remained unknown. Here, we address this gap by characterising type II ELO mutants and show that disruption of FAE remodels the parasite lipidome, impairs fitness under fatty acid-limiting conditions, and reduces virulence and chronic persistence in mice. These findings identify FAE as a key metabolic pathway underpinning adaptation and persistence during chronic infection.

## Introduction

The Apicomplexan *Toxoplasma gondii* is a highly prevalent parasite with a capacity to infect a broad range of organisms (Kim and Weiss, 2004; Molan et al., 2019; Montazeri et al., 2020; Yang et al., 2020; Hajimohammadi et al., 2022). *T. gondii* possesses numerous, and in some cases functionally redundant, metabolic pathways (Xia et al., 2018; Walsh et al., 2022; Rimple et al., 2025) which likely supports its successful parasitism and ability to adapt to varying growth niches.

Fatty acids (FA) are crucial nutrients for *T. gondii* that are obtained by a combination of (i) scavenging from the host cell, and (ii) *de novo* synthesis within parasite cells (Shunmugam et al., 2022). These FA are essential for bulk phospholipid synthesis, membrane biogenesis for daughter cell formation, cell signalling, and lipid droplet storage, all together maintaining intracellular development, propagation and survival (Dass et al., 2021; Chen et al., 2021; Nolan et al., 2018). The constant flux of scavenged FA, also generated by the catabolism of scavenged phospholipids from the host (Dass et al., 2021; Lévêque et al., 2017; Sheokand et al., 2023), must be channelled towards neutral lipid synthesis and stored in parasite lipid droplets. These droplets are specifically mobilized for parasite division, thus enabling timely membrane biogenesis, and prevention of lipotoxicity induced cell death (Dass et al., 2021; Dass et al., 2024). For FA *de novo* synthesis, *T. gondii* possesses two fatty acid synthesis pathways in addition to FA elongases, all together illustrating the complexity of FA metabolism in the parasite **(Figure 1)** (Waller et al., 1998; Ramakrishnan et al., 2012; Tymoshenko et al., 2015; Shunmugam et al., 2022). The parasite bears a cytosolic Type I Fatty acid synthesis pathway (FASI), considered non-essential and likely inactive during its virulent fast-replicating tachyzoite form, the lifecycle stage responsible for acute toxoplasmosis (Tymoshenko et al., 2015). *Toxoplasma* further habours a prokaryotic type II FA synthesis pathway (FASII) in the parasite apicoplast, which is critical for parasite survival as it provides the bulk of *de novo* synthesised FA (Mazumdar et al., 2006; Ramakrishnan et al., 2012; Amiar et al., 2016). The FASII pathway is initiated by the import of the glycolytic intermediate phopsphenolpyruvate (PEP), which is generated by parasite-mediated glycolysis of host scavenged glucose (Lim et al., 2010; Ramakrishnan et al., 2012; Amiar et al., 2016). PEP is then imported into the apicoplast via a plant-like triose phosphate transporter, (*Tg*APT1) (Mullin et al., 2006; DeRocher et al., 2008) and used to generate acetyl-acyl carrier protein (ACP), the substrate of the FASII pathway, via pyruvate kinase II (*Tg*PYKII) and pyruvate dehydrogenase (*Tg*PDH) (Lim et al., 2010; Oppenheim et al., 2014; Xia et al., 2019). The FASII pathway is composed of four core active enzymes allowing generation of FA chains through the addition of two carbons from the acetyl-ACP subtrate per cycle within the pathway. The major products of the FASII are myristic acid (C14:0), and palmitic acid (C16:0)(Ramakrishnan et al., 2012; Amiar et al., 2016) (**Figure 1**).

**Figure 1:**
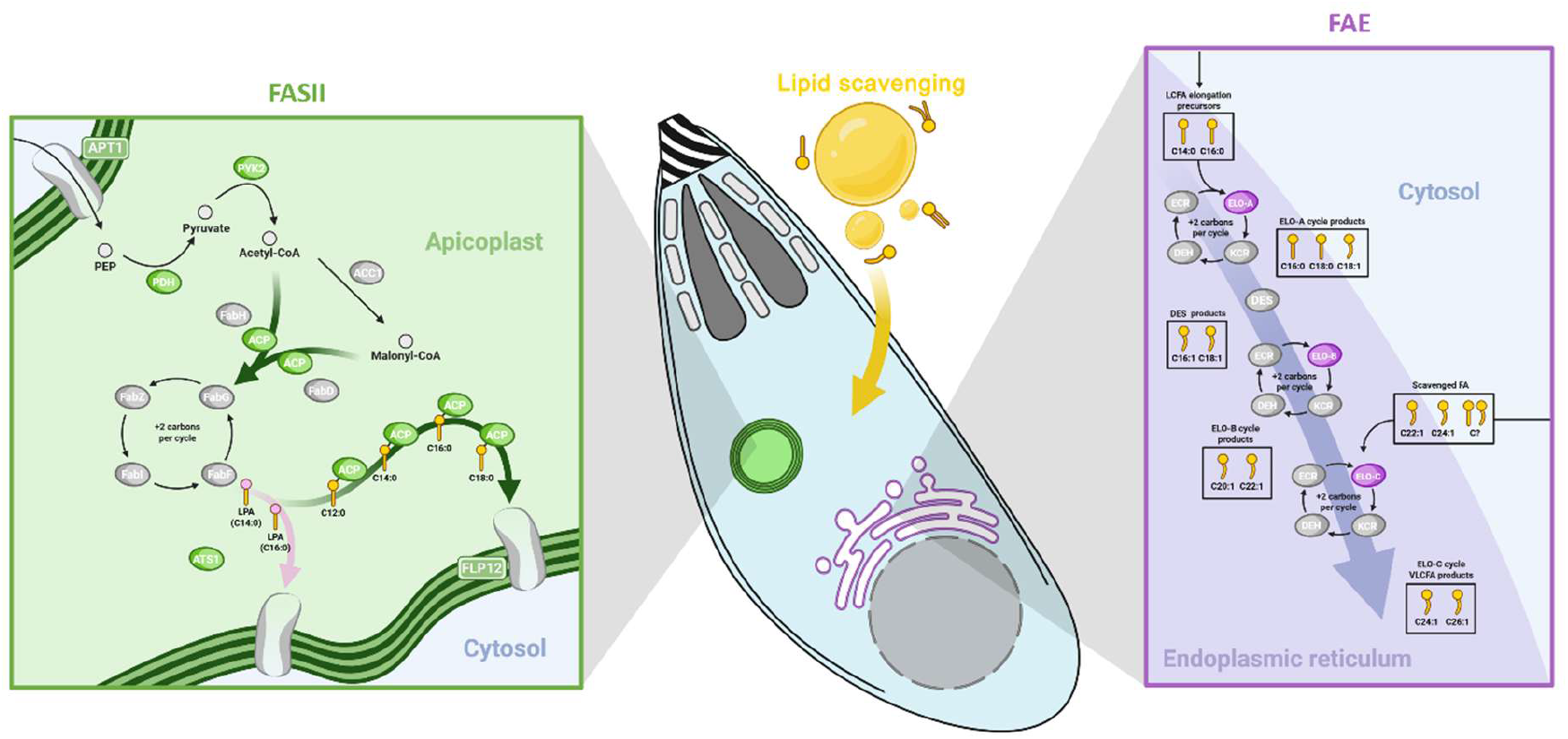
Synthesis of long chain fatty acid species in T. gondii. Highlighted enzymes initiate fatty acid extension or ferry their components. Made with BioRender.com. Abbreviations: Acetyl-CoA carboxylase 1 (ACC1), acyl carrier protein (ACP), apicoplast transporter 1 (APT1), glycerol 3-phosphate acyltransferase 1 (ATS1), coenzyme A (CoA), hydroxyacyl-CoA dehydratase, desaturase (DES), enoyl-CoA reductase (ECR), elongase (ELO), fatty acid biosynthesis (Fab), ketoacyl-CoA reductase (KCR), long chain fatty acid (LCFA), lysophosphatidic acid (LPA), pyruvate dehydrogenase (PDH), phopsphenolpyruvate (PEP), pyruvate kinase II (PYK2), very LCFA (VLCFA).

We previously showed that *T. gondii* can sense the nutritional content of the host and rewire its metabolic program, making some pathways more essential than others under specific nutritional conditions (Botté et al., 2013; Amiar et al., 2020; Bitew et al., 2025). This plasticity promotes parasite adaptation to both favourable and limiting host conditions, maintaining intracellular development and survival. The apicoplast FASII is one such pathway, with its activity dependent on host nutritional content: under high host nutrient content, as mimicked by *in vitro* culture complemented with 10% foetal bovine serum (FBS), the apicoplast FASII activity stays basal and the parasite compensates by massive host lipid scavenging; however under lower nutrient content (eg 0%, 1% FBS), *T. gondii* upregulates FASII activity, which becomes essential for its survival (Botté et al., 2013; Amiar et al., 2020). In these conditions, the FASII can also be artificially bypassed by addition of exogenous FA sources that can be efficiently scavenged and thus partially rescue parasite growth (Krishnan et al., 2020; Liang et al., 2020; Chen et al., 2025). *De novo* synthesised FA from the apicoplast, specifically C14:0 and C16:0, can be incorporated into a lysophosphatidic acid (LPA) precursor via a plant-like acyltransferase in the apicoplast, *Tg*ATS1 (a homolog of *Pf*ApiG3P in *Plasmodium falciparum*) (Amiar *et al*., 2016; Shears *et al*., 2017). Exported to the endoplasmic reticulum (ER), these apicoplast FA are combined with scavenged FA to generate the bulk of phospholipids needed for parasite division (Amiar et al., 2016; Shears et al., 2017). It was proposed that apicoplast synthesised FA such as C14:0 can be directly exported to the parasite ER via a novel essential P5-adensoine triphosphatase (ATPase) transporter conserved among apicoplast-bearing parasitic apicomplexa, named *Tg*FLP12 in *Toxoplasma* (Arnold et al., 2025). Together, apicolast free FA (FFA) and/or FA incroporated in the LPA precursor, reach the three ER located elongases (ELOs) to be elongated further (Ramakrishnan et al., 2012; Ramakrishnan et al., 2015; Dubois et al., 2018; Kloehn et al., 2021).

Triglycerides and multiple phospholipids use long chain fatty acids (LCFA) and very long chain fatty acids (VLCFA) species with chains exceeding 18 and 22 carbon atoms, respectively (Erdbrügger and Fröhlich, 2021). In *T. gondii*, VLCFA are made by the FA elongation (FAE) pathway (Ramakrishnan et al., 2012; Ramakrishnan et al., 2015). Each FAE complex is comprised of a ketoacyl-CoA reductase (KCR), hydroxyacyl-CoA dehydratase (DEH), enoyl-CoA reductase (ECR) and one of three elongase (ELO) homologues **(Figure 1)**. Fuelled by a cytosolic pool of acetyl-CoA and malonyl-CoA (Dubois et al., 2018; Kloehn et al., 2020), *Tg*ELO-A initiates extension of C14:0 and C16:0 into stearic acid (C18:0), and desaturated palmitoleic acid (C16:1) to oleic acid (C18:1) **(Figure 1)** (Ramakrishnan et al., 2012). C18:1 is then channelled by *Tg*ELO-B mediated FAE cycles to erucic acid (C22:1) and followed by *Tg*ELO-C mediated FAE cycles to palmitoleic acid (C26:1) **(Figure 1)** (Ramakrishnan et al., 2012). All *Tg*ELO enzymes efficiently produce unsaturated LCFA and/or VLCFA, however, *Tg*ELO-C differs, as it also uses both scavenged and *de novo* synthesised FA (Ramakrishnan et al., 2015). Depletion of ECR or DEH has been shown to arrest ER associated FAE. While ECR was found essential, reduced DEH activity was rescuable through exogenous supplementation of specific unsaturated LCFA and VLCFA species (Ramakrishnan et al., 2015). However, individual depletion of *Tg*ELO enzymes, barring moderate fluctuations in FA availability, reportedly incurred no growth defect *in vitro* in a type I *T. gondii* strain (Ramakrishnan et al., 2012).

Nonetheless, evidence points to *Tg*ELO’s pivotal role under certain conditions. Downregulation of cholesterol synthesis in interferon gamma (IFN-γ)-activated macrophages has been linked to increased *Tg*ELO-B expression in the type I RH strain of *T. gondii* (Wang et al., 2020) and separately a CRISPR screen flagged *Tg*ELO-B (TGME49_242380) as essential to the same parasite strain during acute mouse infection (Giuliano et al., 2023). It is clear from these studies that FAE provides integral components to the lipidome in various conditions, however their importance in establishing and maintaining a long-term chronic infection has not been tested. No targeted characterisation study of FAE system components *in vivo* has yet been conducted and furthermore, our recent CRISPR-based whole genome screen under high vs low host nutrient content suggested that *Tg*ELOs have differential importance depending on the host nutritional content (Bitew et al., 2025)

To tackle this, we analysed knockouts of *Tg*ELO-A, and both *Tg*ELO-A and *Tg*ELO-B in a cystogenic Type II *T. gondii* strain (ME49). Mutants were phenotypically characterised in FA replete and depleted conditions revealing growth defects linked to parasite replication and invasion. Mass spectroscopy-based lipidomic analyses revealed that both single and double mutants are resilient to environmental FA availability pressures. Likewise, *Tg*ELO depletion incurred virulence reduction in BALB/c mice and greatly reduced brain cyst burdens during chronic infection. Our results join a growing number of studies demonstrating the complex interplay between *de novo* metabolism and scavenging in *Toxoplasma* and highlight the importance of FAE for virulence and persistence.

## Materials and methods

### Cell lines and routine culturing conditions

ME49, ME49ΔELO-A, and ME49ΔELO-AΔELO-B *T. gondii* were co-cultured with human foreskin fibroblasts (HFF)-1 (ATCC, SCRC-1041) in Dulbecco’s modified essential media (DMEM, Gibco, Thermo Fisher Scientific) high glucose, with L-glutamine (11965), supplemented with 10% [v/v] foetal bovine serum (FBS), and 1% [v/v] penicillin/streptomycin (pen/strep) antibiotic mix (Gibco, Thermo Fisher Scientific). Cultures were maintained in a 37°C humidified 5% CO_2_ incubator.

### Construct synthesis

*T. gondii* sequences were accessed and retrieved from ToxoDB (Alvarez-Jarreta et al., 2024). Guide RNA in close proximity to the start and stop codons of *Tg*ELO-A (TGME49_253880) and *Tg*ELO-B (TGME49_242380) were designed using CHOPCHOP (Labun et al., 2019). Guides were annealed and ligated into the CRISPR-Cas9 expressing pG474 vector (Curt-Varesano et al., 2016) as well as concentrated by transformation into, and subsequent extraction from DH5α chemically competent *Escherichia coli*. The pyrimethamine resistance cassette, dihydrofolate reductase (DHFR), with homology flanks for insertion into the *Tg*ELO-A open reading frame (ORF) was amplified from the pDT7S4-G13M5 template plasmid (Sheiner et al., 2011) by polymerase chain reaction (PCR). The green fluorescence cassette, mNeonGreen, with homology flanks for insertion into the *Tg*ELO-B ORF was amplified from the pHL018-pBM014_SAG1-mNeonGreen_TUB1-dTomato template plasmid (gifted by Dr Clare Harding) (Aghabi et al., 2023) by PCR.

### Transfection and cell line generation

ME49 *T. gondii* tachyzoites were transfected with CRISPR-Cas9 vectors targeting upstream and downstream of the gene of interest, and amplified DHFR insertion cassettes by electroporation. Transfectants were selected with 1:1000 pyrimethamine for three days, and a clonal ME49ΔELO-A mutant was isolated. ME49ΔELO-A *T. gondii* tachyzoites were later transfected with dual CRISPR-Cas9 vectors and amplified mNeonGreen insertion cassette by electroporation. After three days, green fluorescent populations were identified by flow automated cell sorting and a clonal ME49ΔELO-AΔELO-B mutant was isolated.

### Cell line validation

Genomic integration of insertion cassettes at GOI loci were initially validated by PCR. Parasite material was pelleted and resuspended in 1:10 20 mg/mL Proteinase K in Tris-EDTA (TE) buffer. Samples were incubated at 60°C for 45 min and then 95°C for 10 min in a thermocycler. PCR reactions were run using GoTaq® Hot Start Green Master Mix and the following programme: Lid was heated to 112°C, polymerase activation 95°C 2 min, then 30 cycles of denaturation 95°C 30 s, elongation 55°C 30 s, and annealing 72°C 1 min/kb. A final elongation cycle of 72°C 10 min was employed to the reaction mixture prior to agarose gel electrophoresis.

For further validation, gene expression was interrogated by reverse transcriptase quantitative-PCR (RT-qPCR). For this, Intracellular parasites were mechanically egressed, filtered through a 0.3 µm membrane to remove host cell debris, pelleted, and RNA was extracted. RNA extraction was conducted using a RNeasy Mini Kit (Qiagen) and RNA concentration normalised to <2 µg via Nanodrop™. Sample RNA was subjected to DNAse treatment and converted to complementary DNA (cDNA) using a High-Capacity RNA-cDNA™ Kit (Applied Biosystems, TFS). RT-qPCR reactions were run using Power Sybr™ Green Master Mix (Applied Biosystems, TFS) and 10 ng cDNA. Sample loaded MicroAmp® Optical 96-Well Reaction Plates (Applied Biosystems, TFS) were run on a 7500 Real Time PCR system (Applied Biosystems, TFS) to the following programme: stage one at 50°C for 2 min and stage two at 95°C for 10 min. Followed by 40 cycles of stage three at 95°C for 15 s, and 60°C for 1 min. And a final stage four at 95°C for 15 s, 60°C for 1 min, 95°C for 15 s, and 60°C for 15 s. Relative expression was calculated via the double CT method (Livak and Schmittgen, 2001; Schmittgen and Livak, 2008).

All oligo sequences used in construct synthesis, PCR, and RT-qPCR are listed in Table 1.

### Plaque assays

Intracellular parasites were mechanically egressed and 500 tachyzoites of each strain seeded onto HFF monolayer coated 6-well plates. Media consisted of DMEM (11965, Gibco, Thermo Fisher Scientific) supplemented with either 10%, 5% or 1% FBS and plates were incubated for seven days under routine conditions. Well contents were fixed with ice cold methanol and stained with 0.4% crystal violet solution. Well contents were imaged and plaque area was quantified using FIJI software (Schindelin et al., 2012). Plaque numbers were counted within the perimeters of two 14mm diameter pre-set boundaries.

### Replication and invasion assays

For replication assays, parasites were subjected to pre-treatment in either media condition: DMEM (1165) supplemented with 10% FBS, or 1% FBS, or glucose deficient DMEM (11966, Gibco, Thermo Fisher Scientific) supplemented with 5% [v/v] FBS and 1% [v/v] pen/strep (Gibco, TFS) for three days. HFF-coated glass coverslips in 24-well plate conformations were pre-treated in respective media conditions for >1 day. Intracellular parasites were mechanically egressed, filtered, seeded onto HFF monolayers, and incubated for 1 hr before a gentle wash with the respective condition media to remove extracellular parasites. Cultures were then incubated for 24 hrs before a gentle phosphate buffered saline (PBS) wash and fixation in 4% Paraformaldehyde (PFA).

For invasion assays, parasites and HFF coated glass coverslips were subjected to FBS-related media condition pre-treatment, as described in the replication assay. After harvesting, parasites were seeded onto HFF monolayers, and incubated for 30 min. Well contents gently washed with PBS to remove unattached extracellular parasites, prior to fixation in 4% PFA.

Cells were blocked with 0.2% Triton X (TX)-100, 2% bovine serum albumin (BSA) in PBS. Coverslip contents were labelled with rabbit α-inner membrane complex (IMC)1 primary antibody (1:1000, gifted by Professor Dominique Soldati-Favre) in blocking buffer, and stained with Goat anti-Rabbit IgG (H+L) Cross-Adsorbed Secondary Antibody, Alexa Fluor™ 488 (1:1000, Invitrogen, TFS). For invasion assays, cells were initially blocked with 2% BSA in PBS, labelled with rat α-surface antigen (SAG)1 primary antibody (1:400, gifted by Prof Vernon B. Carruthers) and stained with Goat anti-Rat IgG (H+L) Cross-Adsorbed Secondary Antibody, Alexa Fluor™ 594 (1:1000, Invitrogen, TFS) in non-permeabilising blocking buffer. Coverslips were washed with Mili-Q water and mounted to glass slides with DAPI Fluormount-G® mounting medium (SouthernBiotech).

### Differentiation assays

Approximately 1.5×10^5^ mechanically egressed tachyzoites were seeded onto HFF-coated glass coverslips within 6-well plate conformations and cultured under routine conditions for 24 hrs. Media was then replaced with differentiation inducing media: Roswell Park Memorial Institute (RPMI) 1640 medium (Sigma-Aldrich, Merck) without NaHCO_3_, supplemented with 50 mM HEPES (Sigma-Aldrich, Merck), 1% [v/v] FBS, 1% [v/v] pen/strep antibiotic mix, and made up to pH 8.3 with NaOH. Differentiation-inducing media was replaced daily and cultures were maintained for seven days in a 37°C humidified 0% CO_2_ incubator, prior to fixation with 10% formalin.

Cells were permeabilised with 0.1% TX-100 in PBS and then blocked with 1% BSA in PBS. Coverslip contents were labelled with rat α-*Tg*SAG1 (1:400) and rabbit α-bradyzoite specific antigen 1 (BAG1, 1:500, gifted by Dr Vernon B. Carruthers) in 0.1% BSA in PBS, and stained with Goat anti-Rat Immunoglobulin G (IgG) (H+L) Cross-Adsorbed Alexa Fluor™ 350 (1:1000, Invitrogen, TFS) and Goat anti-Rabbit IgG (H+L) Cross-Adsorbed Alexa Fluor™ 594 (1:1000, Invitrogen, TFS), as well as Dolichos Biflorus (Horse Gram) Agglutinin (DBA) fluorescein isothiocyanate (FITC, 1:1000, Invitrogen, TFS). Coverslips were washed with Milli-Q water and mounted onto glass slides with ProLong™ Diamond Antifade Mountant (Life Technologies, TFS)

Images were acquired as described below and exported as TIFF files for analysis. GFP fluorescence was used as a proxy for cyst size and quantified in ImageJ using an automated macro adapted from (Di Cristina et al., 2017). Images were converted to 8 bit, spatially calibrated to 0.645 µm per pixel, and thresholded using the Intermodes method to generate binary masks. Following morphological opening, cysts were identified by particle analysis with size limits of 130 to 1,900 µm² and circularity of 0.30 to 1.00, and regions of interest were measured automatically to determine cyst area.

### Fluorescence microscopy

A DeltaVision Core microscope (Applied Precision), Leica DMi8 inverted widefield microscope with LED light source (Leica Microsystems), and ZEISS Axio Observer Z1 inverted microscope were used for image acquisition. Organelle tagging, replication and invasion assay image acquisition parameters consisted of 16-25 stacks of 0.3 µm slices taken at 100x magnification. Differentiation assay image acquisition was conducted at 40x magnification. Image analysis was done using softWoRx (Applied Precision), FIJI (Schindelin *et al*., 2012), and Zen 3.0 blue edition (ZEISS) software.

### Immunofluorescent staining and phenotypic analysis of organelle morphology

Mechanically egressed parasites were seeded onto HFF monolayer coated glass coverslips in 24-well plate format and grown in DMEM (11965) supplemented with 1% FBS, under routine incubation conditions. After 36 hours, media was removed and coverslips gently washed with PBS prior to application of 4% PFA fixative.

Cells were blocked as described in the replication assay with the following changes. Cells were labelled with either rabbit anti-HSP60 (1:1000, gifted by Dr Boris Striepen), rabbit anti-MIC5 (1:1000, gifted by Prof Vernon B. Carruthers), and or mouse anti-ROP7 (1:1000, gifted by Prof Vernon B. Carruthers), as well as stained with either Goat anti-Rabbit IgG (H+L) Cross-Adsorbed Alexa Fluor™ 594 (1:1000, Invitrogen, TFS), Goat anti-Rabbit IgG (H+L) Cross-Adsorbed Secondary Antibody, Alexa Fluor™ 488 (1:1000, Invitrogen, TFS), or Goat anti-mouse Alexa Fluor™ 647 (H+L) Cross-Adsorbed Secondary Antibody (1:1000, Invitrogen, TFS). Coverslips were mounted to glass slides with DAPI Fluormount-G® or Fluormount-G® mounting medium (SouthernBiotech).

Images were acquired as previously described and exported in TIFF format for analysis. Raw images were deconvolved with ™Huygens Essential software using default wizard guided settings. 50 vacuoles per biological replicate (n=3) were counted and either nuclear, apicoplast, microneme or rhoptry morphology assessed. Morphs were categorised as ‘normal’ based on previous descriptions in the literature (Breinich et al., 2009; Sheiner et al., 2010; Jacot et al., 2013; Kremer et al., 2013; Dogga et al., 2017; Venugopal et al., 2017; Morlon-Guyot et al., 2018; Ben Chaabene et al., 2024; Pasquarelli et al., 2024; Tagoe et al., 2024), and ‘abnormal’ when deviating or not visible.

### Transmission-electron microscopy

Parasites were seeded onto 10 cm dishes containing HFF lawns and grown in DMEM (11965) supplemented with 1% FBS, under routine incubation conditions. After three days, when parasite vacuoles were confluent, cells were washed with PBS and fixed in 2.5% glutaraldehyde, 4% PFA in 0.1 M phosphate buffer. After fixing, samples were serially washed 0.1 M phosphate buffer, and then post-fixed in 1% osmium tetroxide, 1.25% potassium ferrocyanide in 0.1 M phosphate buffer [v/v] for 1 hr in darkness. Samples were then washed with distilled water and contrasted together with 0.5% aqueous uranyl acetate for 1 hr in darkness. Samples were subjected to acetone mediated dehydration in increasing concentrations (30%, 50%, 70%, 90%, 100%) and embedded in epoxy resin. 50 nm ultra-thin sections were cut using a Leica Ultramicrotome (Leica Microsystems), collected onto 100 mesh grids covered with formvar and contrasted with 2% aqueous uranyl acetate. Sections were imaged using a JEM-1400Flash transmission electron microscope (Jeol) operating at 80 kV.

### Lipidomic profiling

Parasites were heavily seeded into 175 cm flasks containing HFF lawns and cultured in DMEM (11965) supplemented with 10% or 1% FBS. Following 72-96 hrs incubation, whereby parasite vacuoles were confluent, both intracellular and extracellular parasites were quickly harvested and metabolically quenched by rapid chilling to 0°C. Quenched samples were maintained at 4°C and washed with ice cold PBS.

Lipid extraction and gas chromatography-mass spectrometry (GC-MS) analysis was as previously described (Dubois et al., 2018) using a GCMS (Agilent7890B gas chromatograph and 5977A Series mass selective detector). Individual fatty acid abundance was quantified via Mass Hunter quantification software (Agilent) and normalised through the internal standard and cell number.

### Stable isotope metabolite labelling experiment

HFF monolayers grown to confluency in 175 cm flasks were heavily seeded with *T. gondii* tachyzoites and cultured in glucose deficient DMEM (11966) supplemented with 10% [v/v] FBS, 1% [v/v] pen/strep antibiotic mix, and 8 mM of either U^12^-C glucose or U^13^-C glucose. Cultures were incubated, processed, and extracted lipids analysed as previously described for lipidomic profiling. After correcting for concentration of U^13^-C glucose within the culture media, ^13^C incorporation for each FA species was calculated as previously described (Dass et al., 2021)

### Mouse maintenance

All animal procedures were pre-approved by the Institutional Ethics Committee of the Institut Pasteur de Montevideo (CEUA Nos. 019-19 and 023-22). Bagg Albino c (BALB/c) mice, bred *in house* were maintained in a dedicated facility at the Laboratory Animals Biotechnology Unit, Institut Pasteur de Montevideo, underspecific pathogen-free conditions and monitored daily. Under routine and experimental conditions, groups of five mice were maintained in individually ventilated cage racks (1285 L, Tecniplast) at 19-21°C, 30–70% relative humidity, under negative pressure during biocontainment, and with a 14 hrs light 10 hrs dark cycle.

Humane endpoints were strictly adhered to per the established norms. Euthanasia was performed by cervical dislocation under deep-sleep anaesthesia. Mice were injected intraperitoneally with freshly prepared 10 mL/kg ketamine–xylazine 2% solution, corresponding to 110 mg/kg ketamine (phs® Pharmaservice Montevideo-Uruguay) and 13 mg/kg xylazine (Laboratorios Microsules Uruguay S.A.). The absence of motor response to nociceptive stimulation of the forelimbs and hindlimbs was used to confirmed depth of anaesthesia. All experiments adhered to the relevant guidelines outlined in the animal experimentation protocol, abiding by the national animal protection law (No. 18.611) and the ARRIVE guidelines (Percie du Sert et al., 2020).

### Mouse survival experiment

For survival experiments, 6–8-week-old male mice were inoculated intraperitoneally with 50 or 1000 tachyzoites of either ME49 *T. gondii* or derived mutants and their health monitored over the course of infection. A score sheet was used to record changes in the appearance, physical condition, and behaviour of infected mice.Animals expressing the following symptoms were euthanised by cervical dislocation: pronounced inactivity/lack of movement, pronounced hunched posture, pronounced ataxia, abnormal breathing (rapid or laboured), low food or water intake and/or weight loss exceeding 20% initial weight.

### Mouse brain cyst burden assessment

For brain cyst burden assessment, brains from chronically infected mice were harvested 30 days post-infection (dpi) and homogenised using a tissue grinder. 250 µL brain homogenate samples were pelleted and resuspended in 3% formalin fixative in PBS, then pelleted and resuspended in 0.1 mM glycine in PBS, then pelleted and resuspended in blocking buffer: 3% BSA, 0.2% Triton X-100 in PBS. Samples were resuspended and stained with DBA, Fluorescein (1:500) in blocking buffer, prior to three washes with 0.2% Triton X-100 in PBS. After loading onto a glass slide, the number of cysts in each sample was counted via fluorescence microscopy.

## Results

### Sequential deletion of TgELO-A and TgELO-B yields viable ME49 *T. gondii*

To explore the role of FAE in cytogenic *T.* gondii, we generated knockouts of the two first *Tg*ELOs in the FAE pathway in the type II ME49 strain. Simultaneous depletion of *Tg*ELO-A and *Tg*ELO-B was previously suggested as refractory in the type I RH *T. gondii* strain (Ramakrishnan et al., 2012), therefore we opted for a sequential deletion approach. Knockout of *Tg*ELO-A was generated through use of a pyrimethamine resistance selection cassette, whereas sequential knockout of *Tg*ELO-B in ME49ΔELO-A was generated using a green fluorescence reporter insertion cassette **(Figure 2A)**. PCR analysis confirmed loss of *Tg*ELO exons and cassette integration into each respective open reading frame **(Figure 2B)**. Depletion of *Tg*ELO-A and *Tg*ELO-B mRNA transcripts was confirmed via RT-qPCR **(Figure 2C)**. These data validate that *Tg*ELO-A and *Tg*ELO-B were both dispensable for *T. gondii* ME49 growth in high nutrient content (10% FBS) culture conditions.

**Figure 2:**
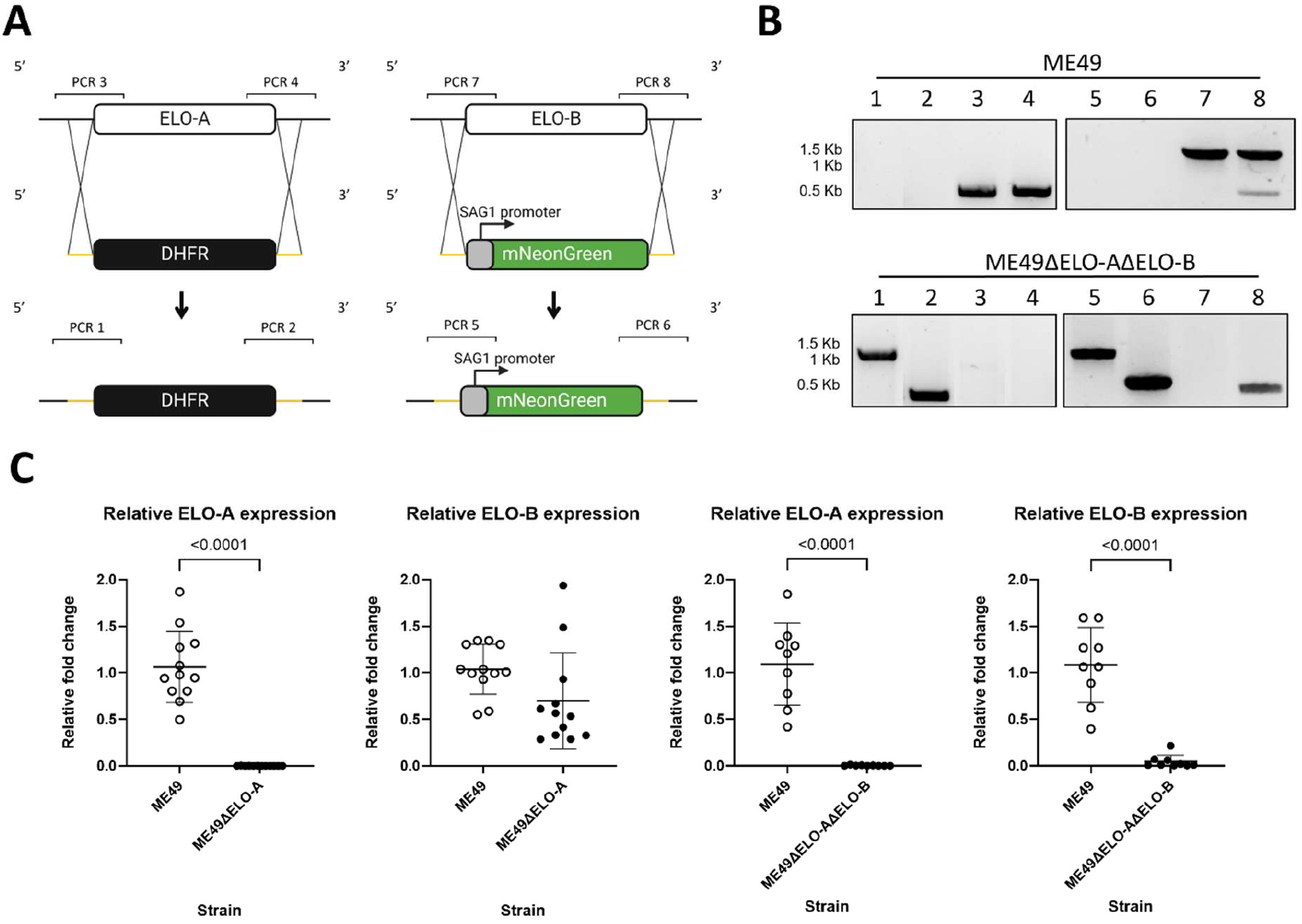
Knockout mutant generation. **(A)** Strategy for complete CRISPR-Cas9 mediated gene knockout using DHFR resistance and mNeonGreen fluorescence selection cassettes with validation by PCR. **(B)** PCR validation of gene of interest deletion in ME49ΔELO-AΔELO-B with ME49 *T. gondii* as a control. Expected amplicon sizes for PCR 1 (primers i + ix) is 1750 bp, PCR 2 (primers iii + x) is 431 bp, PCR 3 (primers i + ii) is 702 bp, PCR 4 (primers iii + iv) is 805 bp, PCR 5 (primers v + xi) is 1318 bp, PCR 6 (primers vii +xii) is 852 bp, PCR 7 (primers v + vi) is 1499 bp, and PCR 8 (primers vii + viii) is 804 bp. **(C)** Fold change in gene of interest mRNA expression relative to the housekeeping gene *Tg*Actin-1 in ME49 strain *T. gondii* and derived mutants. Three individual samples from separate parasite populations per strain were subjected to three-four RT-qPCR runs consisting of three technical repeats. Statistical analysis was performed using independent two tailed t-tests. Error bars represent mean ± SD.

### The *T*. *gondii* lipidome adapts to loss of *Tg*ELO-A and *Tg*ELO-B

ELO enzymes are responsible for initiating reaction cycles in the production of LCFA and VLCFA. Hence, we wondered how production of LCFA and VLCFA was affected in our single and double mutants. To gauge this, we conducted mass spectrometry-based lipidomic analyses and determined the FA profiling in the mutants under replete and depleted nutritional culture conditions. Indeed, we and others have shown that *T. gondii* is capable of adapting to the growth media nutritional content, and in response, can metabolically rewire and adapt to maintain intracellular propagation (Charital et al., 2024a; Charital et al., 2024b; Dass et al., 2021; Bitew et al., 2025). This can also be seen as enzymes essentiality or dispensability under specific nutritional conditions, especially given by the amount of lipids provided by FBS in the culture media (Amiar et al., 2020; Krishnan et al., 2020; Dass et al., 2021; Primo et al., 2021; Bitew et al., 2025).

Under high nutrient content at 10% FBS, we measured a significant increase in smaller saturated and unsaturated LCFA abundance, including: C14:0, C16:0, and C16:1, which are the typical apicoplast FASII products, and the substrates for *Tg*ELOs in both ME49ΔELO-A (P < 0.05 – 0.0001) and ME49ΔELO-AΔELO-B (P < 0.0001) mutants relative to the parental strain **(Figure 3A)**. On the other hand, Larger LCFA species relative abundance including: C18:0, cis C18:1, and eicosenoic acid (C20:1), which are the typical products of *Tg*ELOs (Dubois et al., 2018; Kloehn et al., 2021) significantly decreased (P < 0.0001) in both *Tg*ELO deleted mutants, correlating previous reports in Type I *T. gondii Tg*ELOs **(Figure 3A).** Interestingly, there was a slight, yet significant increase in arachidonic acid (C20:4) relative abundance (P < 0.01) in ME49ΔELO-AΔELO-B compared to ME49ΔELO-A, but not to the parental **(Figure 3A)**. Since the parasite is heterotrophic for arachidonic acid, this might suggest an increase of host FA scavenging by the parasite in the absence of *Tg*ELO-A and *Tg*ELO-B (Bitew et al., 2025).

**Figure 3:**
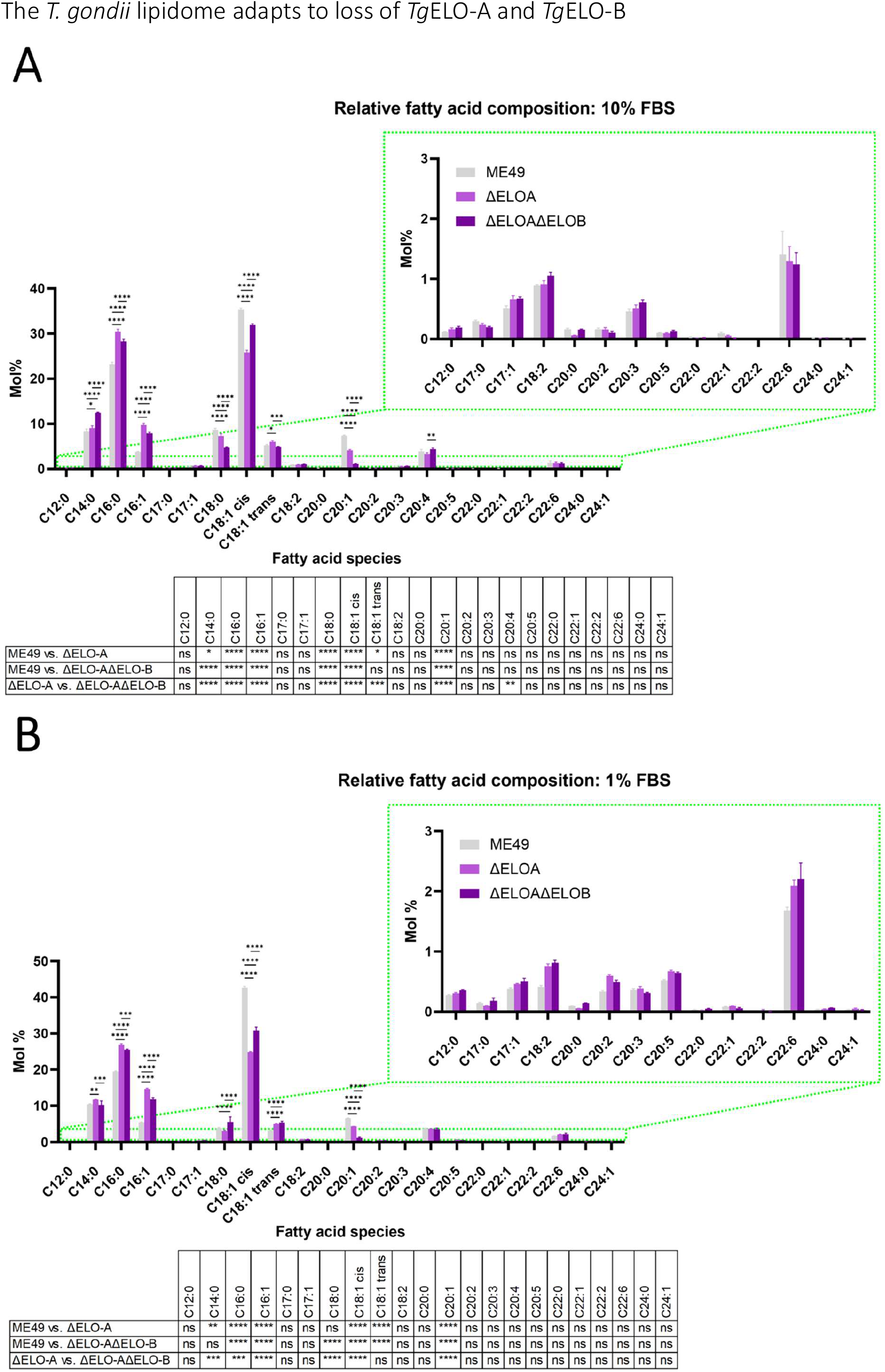
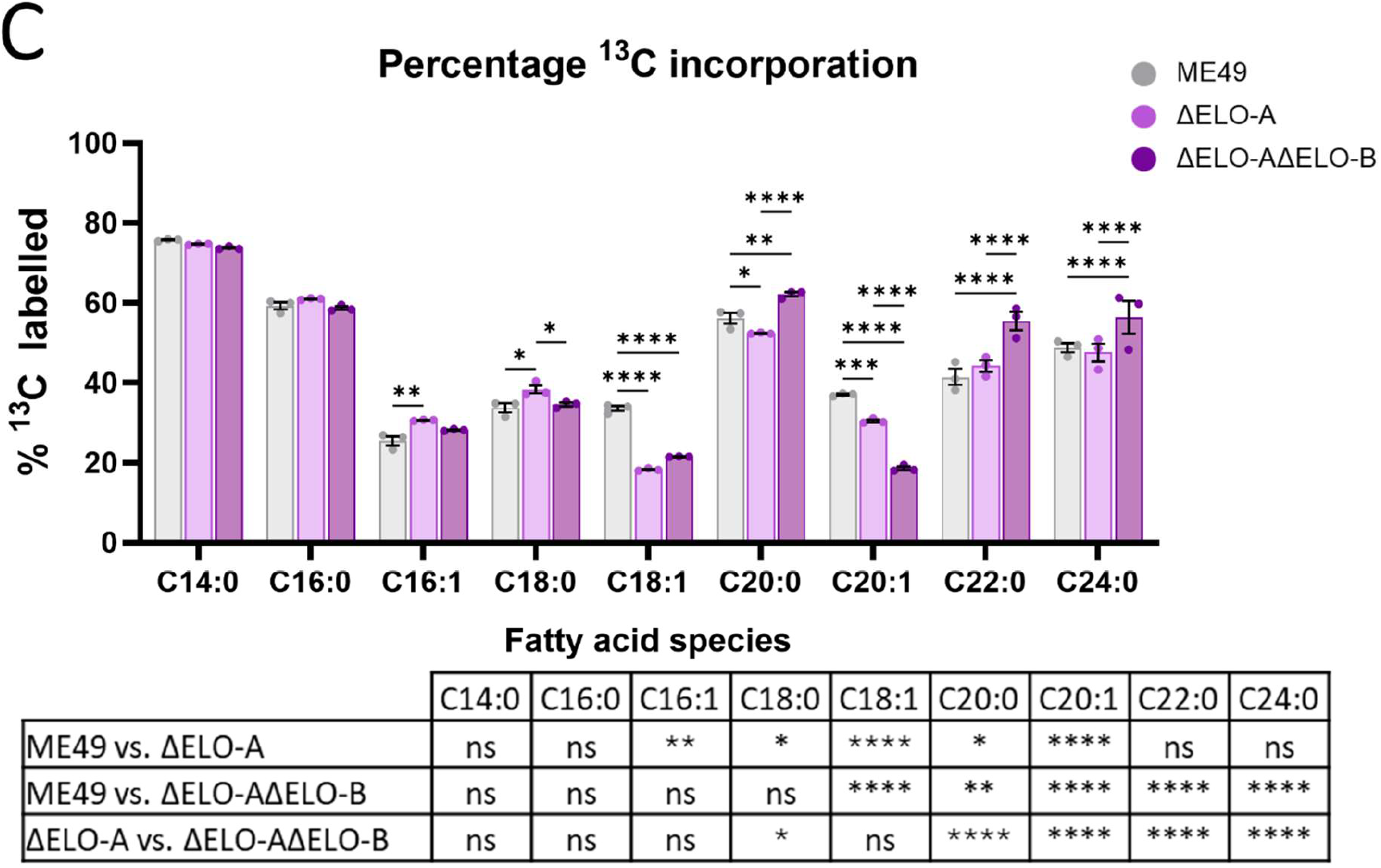
Lipidomic analyses and metabolite labelling experiment. Relative FA composition in ME49 strain *T. gondii* and derived *Tg*ELO depleted mutants in percentage molarity (Mol %). Parasites were cultured in DMEM (11966) supplemented with **(A)** 10% FBS or **(B)** 1% FBS and incubated under routine culturing conditions until confluent. **(C)** Levels of ^13^C derived from U^13^C-glucose incorporation in LCFA and VLCFA species extracted from *Tg*ELO KO and ME49 WT strains, supplemented with 10% FBS, and harvested when confluent. Intracellular and extracellular parasites were quenched at 0°C, prior to processing for gas chromatography-mass spectroscopy (GC-MS). Analysis via Two-way ANOVA with Fishers LSD, n = 3. Unpaired t-test values: * P < 0.05, ** P < 0.01, *** P < 0.001, **** P < 0.0001, not significant (ns).

Culturing *Tg*ELO deleted mutants in a low nutrient content, at 1% FBS, did not result in any further differences in relative FA composition as to those observed at 10%. Likewise our results showed significant increases of C14:0, C16:0, C16:1 (products of FASII and substrates for *Tg*ELOs) and decreases in C18:1, C20:2 (products of *Tg*ELOs). Surprisingly, there was a small but significant increase (P < 0.0001) in C18:0 abundance for ME49ΔELO-AΔELO-B compared to both the parental ME49 and ME49ΔELO-A **(Figure 3B)**. We were unable to detect significant differences in the relative abundance of VLCFA including C22:1, the proposed main product of *Tg*ELO-B, nor detect C26:1, the main product of *Tg*ELO-C. Our findings provide tertiary validation of targeted *Tg*ELO ablation, and are congruent with previous studies of *Toxoplasma* FAE (Ramakrishnan et al., 2012; Ramakrishnan et al., 2015).

To determine the origin of FA elongated by the *Tg*ELOs in type II *T. gondii*, we performed standard stable isotope labelling followed by mass spectrometry-based lipidomic analysis whereby the parasite is grown in the presence of U^13^C-glucose. This allows specific labelling of FA made via the apicoplast FASII, as previously reported (Ramakrishnan et al., 2012; Amiar et al., 2016; Dass et al., 2021). We conducted such approach in either *Tg*ELO-A, or both *Tg*ELO-A and *Tg*ELO-B deficient mutants **(Figure 3C)**. As expected, there were no significant differences in the ^13^C incorporation for C14:0, and C16:0 in the mutants as synthesis occurs via the unmodified apicoplast FASII **(Figure 3C).** However, both mutants displayed significantly lower ^13^C incorporation in C18:1 than parental ME49 *T. gondii* reflecting the absence of *Tg*ELO-A activity, typically generating C18:1 from C16:1 (Ramakrishnan et al., 2012; Dubois et al., 2018)**(Figure 3C)**. Likewise, ME49ΔELO-A had significantly lower (P < 0.001) ^13^C incorporation in C20:1 than ME49 *T. gondii*, reflective of the activity of *Tg*ELO-A and B **(Figure 3C)**. Closer analysis of the isotopologue profile for each of these FA **(Supplementary figure 1A)** confirmed that the FASII produces FA chains up to C14:0 and C16:0 using carbons from glucose, and that both *Tg*ELOs extend these apicoplast FA.

There was a small but significant increase (P < 0.01) in the percentage of ^13^C incorporation in C16:1 for ME49ΔELO-A but not ME49ΔELO-AΔELO-B **(Figure 3C).** In addition, ME49ΔELO-A was found to have a small but significant increase (P < 0.05) in C18:0 compared to the parental ME49 strain and ME49ΔELO-AΔELO-B **(Figure 3C)**. This can be likely explained as C16:1 accumulation due to cessation of its elongation, whereas C18:0 can either be made by the FASII and/or directly scavenged from the host, likely accumulating due to relative lack of other elongated FA. Surprisingly, there were significant increases (P < 0.0001) in ^13^C incorporation for docosanoic acid (C22:0) and lignoceric acid (C24:0) in ME49ΔELO-AΔELO-B compared to the parental ME49 strain **(Figure 3C)**. These findings suggest glucose plays an increasing role in the generation of some but not all LCFA and VLCFA, despite the metabolic obstruction intended to disrupt the mobilisation of apicoplast FASII products to FAE.

### *Tg*ELO deletion induces serum dependent growth defects in tachyzoites *in vitro*

As an obligate intracellular parasite, the *T. gondii* lytic cycle consists of migration, host-cell invasion, replication, and egress before infecting new host cells (Blader et al., 2015). *In vitro*, this lytic cycle generates plaques in a host cell monolayer. Previously, individual depletion of *Tg*ELO-A subunits in the Type I RH strain did not affect *T. gondii* intracellular development under high nutritional content (10% FBS), and according to their phenotypic scores determined in similar conditions (Ramakrishnan et al., 2012; Bitew et al., 2025). However, nothing was known on their putative roles in type II parasites, and/or under different host nutrient environments. Having observed the changes in parasite lipidome, we sought to determine the importance of *Tg*ELOs under varying nutrient environments including lipid deprivation through FBS titration, as recently reported (Bitew et al., 2025). We first conducted plaque assays **(Figure 4A)**. There was no significant difference in plaque numbers observed between wildtype (WT) ME49 and *Tg*ELO KO mutants under high and low nutrient/lipid content at 10% and 1% FBS **(Figure 4B)**. However, in all nutrient conditions (1, 5, 10% FBS) the plaque area made by *Tg*ELO depleted mutants was significantly lower (P ≤ 0.0009) than that of the parental strain, suggesting a slower growth **(Figure 4C)**. As expected, this phenotype was aggravated by decreasing the concentration of FBS **(Figure 4C)**, supporting the role of *Tg*ELOs in metabolic adaptation. Intriguingly, the area of plaques generated by ME49ΔELO-AΔELO-B was significantly smaller than those generated by ME49ΔELO-A (P = 0.0062) in the 10% FBS condition, whereas in the 1% FBS condition the trend was inverted **(Figure 4C)**. This may indicate different roles for *Tg*ELO-B and *Tg*ELO-A in different nutrient content. Comparing plaque area within strains over the FBS titration provided additional support for the link between parasite fitness and culture environment, as all strains showed a significant decrease (P < 0.0001) in plaque area when cultured in 1% FBS compared to 10% **(Figure 4D)**.

**Figure 4:**
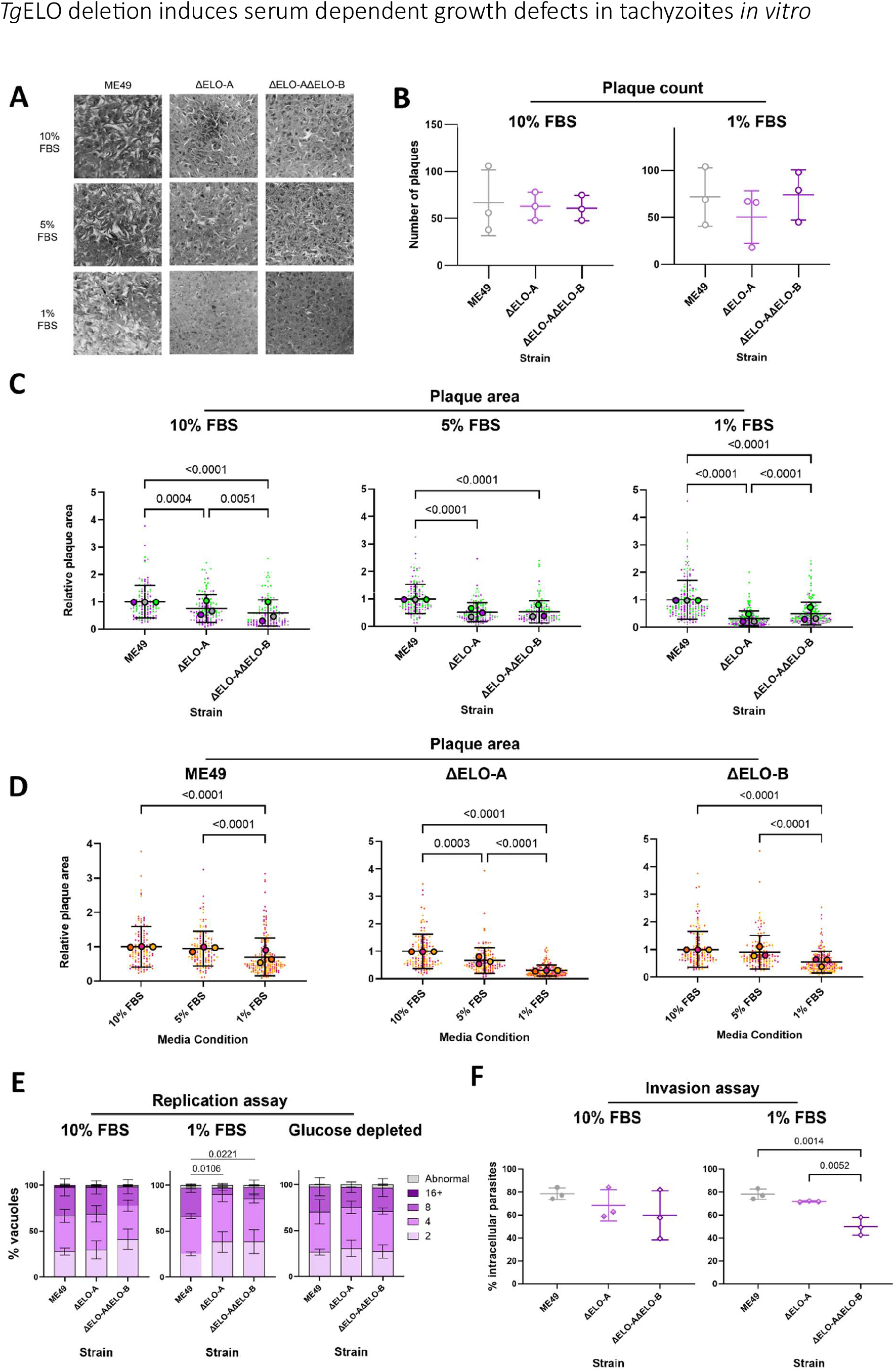
Phenotypic *in vitro* characterisation of ME49ΔELO-A and ME49ΔELO-AΔELO-B tachyzoites. **(A)** Plaque assays comparing growth of ME49 *T. gondii* and derived *Tg*ELO depleted mutants. Approx. 500 mechanically egressed parasites were seeded into HFF monolayers within 6-well plates. Cells were cultured with DMEM (11965) supplemented with 10%, 5%, or 1% FBS, and incubated under routine conditions constituting a humidified 37 °C, 5% CO_2_ incubator for 7 days. Well contents were then fixed and stained with crystal violet. **(B)** Plaque counts were conducted within two areas equalling 14mm diameter. Analysis via one-way analysis of variance (ANOVA) with Tukey’s post hoc. Error bars are mean ± SD, representative of three biological replicates. **(C)** Plaque area relative to ME49 *T. gondii* and **(D)** plaque area of individual *T. gondii* strains relative to the 10% FBS condition, n ≥ 130. Analysis via Kruskal Wallace one-way ANOVA with Dunn’s post hoc. Error bars are mean ± SD representative of three biological replicates. **(E)** Replication rate of ME49 *T. gondii* and derived *Tg*ELO depleted mutants over 24 hrs in different media conditions. Mechanically egressed parasites were seeded onto HFF coated glass coverslips 24-well plates and cultured in DMEM (11965) supplemented with 10% FBS, 1 FBS% or glucose deficient DMEM (11966) supplemented with 5% FBS, in routine incubation conditions. Vacuoles containing a number of parasites not following the 2^n^ rule were categorised as ‘abnormal’. Analysis via one-way ANOVA with Tukey’s post hoc. Error bars are Mean ± SD, representative of three biological replicates, where n = 150. **(F)** Red-green invasion assay of ME49 *T. gondii* and derived *Tg*ELO depleted mutants over 30 min in serum replete and depleted media conditions. Mechanically egressed parasites were seeded onto HFF coated glass coverslips 24-well plates and cultured in DMEM (11965) supplemented with 10% or 1 FBS% and incubated under routine conditions. Analysis via one-way ANOVA with Tukey’s post hoc. Error bars are Mean ± SD, representative of three biological replicates consisting of two technical replicates, where n ≥ 100.

Changes in the area of plaques typically result from defects in stages of the lytic cycle (Srivastava et al., 2020). Given the structural role of LCFA and VLCFA in phospholipid membranes, critical for cell division and motility (Ren et al., 2020; Chen et al., 2021; Erdbrügger and Fröhlich, 2021), we characterised parasite *in vitro* replication and invasion by immunofluorescent staining under physiological levels of glucose and low levels of FBS, together mimicking common physiological conditions in the human patient. We theorised that restricting glucose would introduce further nutritional pressures to lipid synthesis in our *Tg*ELO depleted mutants.

For all mutants tested there was no significant decrease in the replicative rate observed under 10% FBS or glucose depleted media conditions **(Figure 4E)**. There was, however, a significant decrease in the number of larger vacuoles containing more parasites (8 parasites/vacuole) in ME49ΔELO-A (P = 0.0019) and ME49ΔELO-AΔELO-B (P = 0.0061) grown in the 1% FBS media condition **(Figure 4E)**. Likewise, no significant difference in the percentage of successfully invaded parasites between WT and mutants cultured in the 10% FBS condition was observed. However, the mean percentage of successfully invaded parasites significantly decreased by 28.21% (P = 0.0014) for ME49ΔELO-AΔELO-B compared to ME49 when grown at 1% FBS **(Figure 4F)**. These observations suggested the TgELOs deletion and the resulting changes in fatty acid composition affect replication and invasion.

FA are also required during organelle biogenesis, thus we sought to determine whether organelle biogenesis was perturbed under serum depletion. Immunofluorescent staining was performed on parasites grown for 36hrs under 1% FBS. Four organelles critical for replication (nucleus and apicoplast) and invasion (micronemes and rhoptries) were examined via confocal microscopy, with morphology qualitatively assessed as normal or abnormal **(Supplementary figure 3A – B).** Overall, we could not detect a dramatic defect on any of those organelles. While a significant defect in apicoplast morphology was identified in ME49ΔELO-A (P = 0.0458), this was subtle and not seen in ME49ΔELO-AΔELO-B **(Supplementary figure 3C)** suggesting it may not be meaningful. Ultrastructural analysis of all *T. gondii* strains cultured in the 1% FBS condition for 72 hrs was also conducted by transmission electron microscopy to evaluate more organelles. However, here too, no clear ultrastructural defect was apparent **(Supplementary figure 4)**. Therefore, the defects observed in invasion and replication upon TgELOs deletion grown in depleted FBS conditions are not attributed to an organellar defect that manifests in morphological differences.

### *Tg*ELO-B is important for virulence and the establishment of chronic infection

Type II lineage *Toxoplasma* have greater propensity to differentiate between tachyzoite and bradyzoite stages than the type I strains commonly used in *T. gondii* research. As bradyzoites are responsible for chronic infection and transmission, our model grants the opportunity to assess the contribution of FAE to early-stage differentiation *in vitro* and to virulence *in vivo*. Bradyzoite formation is commonly induced *in vitro* via an alkaline stress response (Sanchez et al., 2023; Smith et al., 2021; Tomita et al., 2013). Early differentiation is indicated by the expression of cyst wall proteins, such as *Tg*CST1, and increase in the expression of bradyzoites markers such as *Tg*BAG1 (Bohne et al., 1995) with the concomitant decrease in the expression of tachyzoites markers such as *Tg*SAG1 (Manger et al., 1998; Tomita et al., 2013)**(Figure 5A)**. Loss of *Tg*ELO-A and *Tg*ELO-B had no significant effect on tachyzoite to bradyzoite differentiation *in vitro* with similar stage specific marker expression observed between mutants and WT ME49 *T. gondii* **(Figure 5B)**. Likewise, no significant difference in cyst size *in vitro* was identified **(Figure 5C)**, together indicating the TgELOs are dispensable for *in vitro* differentiation.

**Figure 5:**
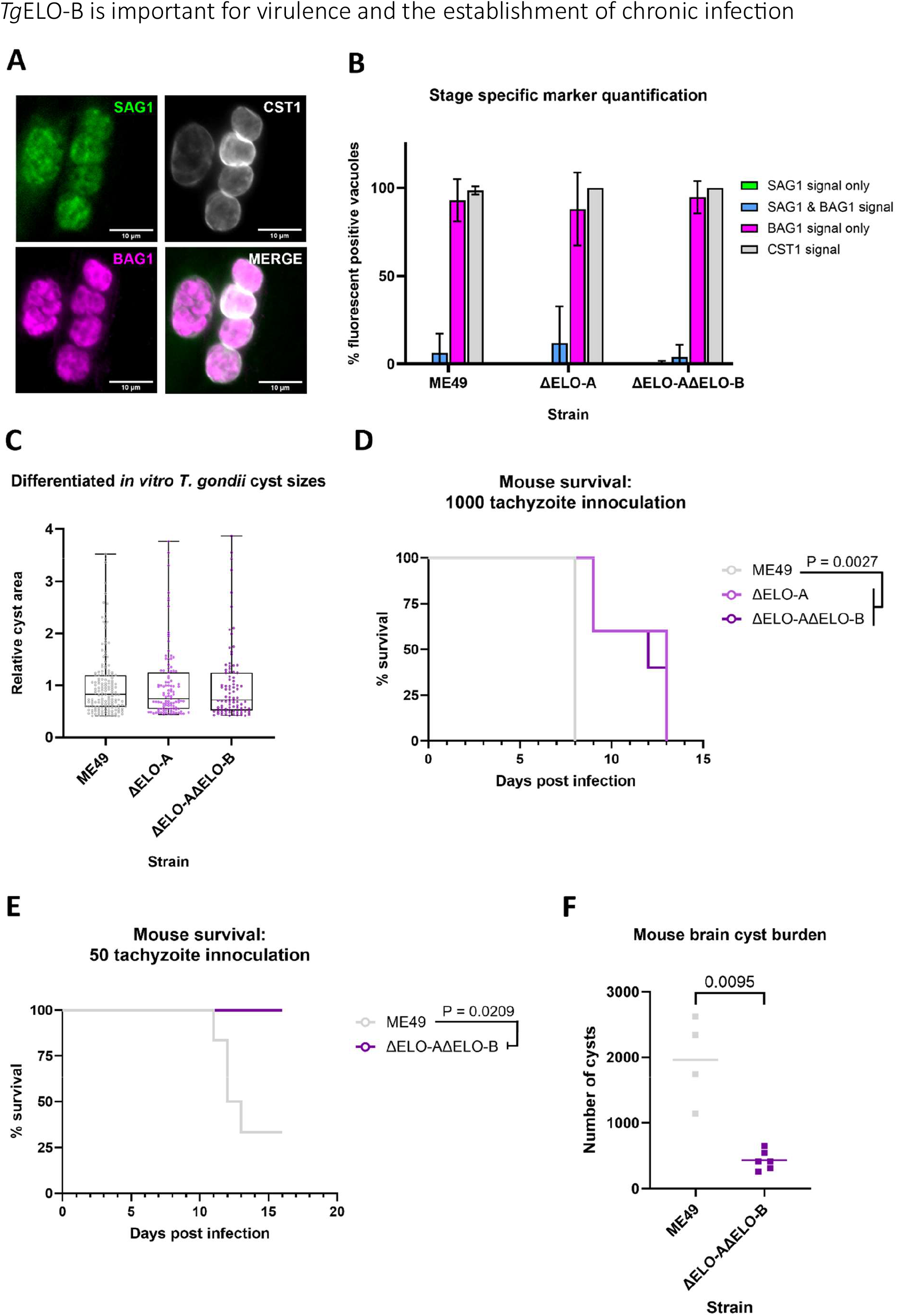
Mouse survival experiment and brain cyst burden assessment. **(A)** Tachyzoite and bradyzoite stage specific marker expression in ME49 *T. gondii* after seven days of culturing in alkaline media. *Tg*SAG1 and *Tg*BAG1 are tachyzoite and bradyzoite markers respectively whilst *Tg*CST1 is a cyst wall marker. **(B)** Differentiation assay quantification of tachyzoite and bradyzoite stage specific markers between ME49 strain *T. gondii* and *Tg*ELO depleted mutants. Parasites were seeded into HFF monolayers within 6-well plates, and after 24 hrs of culture under routine media and incubation conditions, media was replaced with RPMI 1640 supplemented with 1% FBS and made up to pH 8.3. Cells were cultured in a humidified 37°C, 0% CO_2_ incubator seven days and alkaline media replaced daily. Analysis via one-way ANOVA with Tukey’s post hoc. Error bars are Mean ± SD, representative of three biological replicates, n ≥ 25. **(C)** Quantification of parasite cyst area relative to ME49 *T. gondii.* Analysis via Kruskal Wallace one-way ANOVA with Dunn’s post hoc. Boxplot whiskers represent minimum and maximum data points, representative of three biological replicates. Mouse survival curve for infection with ME49, ME49ΔELO-A, or ME49ΔELO-AΔELO-B *T. gondii*. Bagg Albino c (BALB/c) lab mice were inoculated with 1000 **(D)**, or 50 **(E)** tachyzoites per mouse of either parental strain or derived *Tg*ELO depleted mutant *T. gondii* and monitored up to 16 days. Analysis was conducted by Gehan–Breslow–Wilcoxon test (n = 5 for the 1000 tachyzoite dose; n=6 for the 50 tachyzoite dose). **(F)** Surviving mice were sacrificed at experiment endpoint and brain cyst burden determined. Analysis via Mann-Whitney test.

While our *in vitro* growth analyses indicated that both ME49ΔELO-A and ME49ΔELO-AΔELO-B display mild growth defects with respect to their parental strain, these did not prevent continuous culture in any media condition tested. In an *in vivo* CRISPR screen using type I RH *T. gondii Tg*ELO-B (TGGT1_242380) was found to be required for parasite survival (Giuliano et al., 2023). In addition, growth defects become more profound in our TgELO mutants under nutrient depletion. Hence, we hypothesized that TgELOs and the corresponding FAE they make might play a role in enabling parasite survival *in vivo* where they encounter different growth conditions and pressures. To test this, we challenged BALB/c mice with parasite doses similar to previously published LD_50_ for ME49 (Li et al., 2018; Wang and Sibley, 2020; Salman et al., 2021). Mice inoculated with either mutant had significantly increased survival duration (P = 0.0027) by one to five days compared to mice infected with ME49 **(Figure 5D)**. Reducing parasite inoculum enhanced mouse survival, enabling analysis of the transition into the chronic phase **(Figure 5E)**. In these mice, we determined a significant decrease (P = 0.0095) in the brain cyst burdens upon infection with ME49ΔELO-AΔELO-B compared to parental ME49 **(Figure 5F)**, indicating *Tg*ELOs are important for mounting a successful chronic infection in the host.

## Discussion

Studies of the *T. gondii* FAE components in type I strains have revealed their contribution to the production of LCFA and VLCFA *in vitro* (Ramakrishnan et al., 2012; Ramakrishnan et al., 2015). However, while the cystogenic type II lineage is responsible for most toxoplasmosis cases worldwide (Hosseini et al., 2018; Fernández-Escobar et al., 2022), no studies have yet addressed how FAE contributes to the survival of the immune-evasive bradyzoites that are dependent on scavenging for persistence. We sought to tackle this gap through conditional culture serum deprivation and *in vivo* studies to better understand the fitness and metabolic contribution of *T. gondii’s* FAE in this important strain.

Previous attempts to simultaneously deplete *Tg*ELO-A and *Tg*ELO-B were unsuccessful (Ramakrishnan et al., 2012). While raising the possibility that these individually inessential enzymes form a lethal synthetic pair, this was not confirmed. Given that prior iΔDEH mutants were rescued by exogenous LCFA and VLCFA supplementation (Ramakrishnan et al., 2015), we reasoned that elevating the serum concentration in culture could support the survival of ME49ΔELO-AΔELO-B *T. gondii* mutants through *Tg*ELO-C activity. In our study we successfully generated single and double KO of *Tg*ELO and *Tg*ELO-B in the type II ME49 *T. gondii* strain, and validated these mutants at the genetic, transcriptional, and lipidomic level **(Figures 2, 3)**. Though *Tg*ELO-A individually, and *Tg*ELO-A and *Tg*ELO-B simultaneously were found inessential to ME49 *T. gondii in vitro*, both deletional mutants incurred significant observable fitness defects when grown in serum replete conditions **(Figure 4A)**. Plaque assays further indicated that ME49ΔELO-A and ME49ΔELO-AΔELO-B were sensitive to serum depletion, a finding supported by a significant replication defect observed in these mutants, and a host-cell invasion defect in the double knockout, both when cultured in 1% FBS supplemented medium **(Figure 4E-F)**. Our study also corroborates our recent CRISPR screen, demonstrating *Tg*ELO-B influenced *T. gondii* survival under serum depletion *in vitro* (Bitew et al., 2025). While the experiments are performed with different *T. gondii* strains, we suggest that in addition, the growth in 1% foetal calf serum (Ramakrishnan et al., 2012), versus 10% FBS in our study, likely contributed to the different study outcomes.

During host-cell invasion, *T. gondii* uses dedicated secretory organelles to mediate attachment and invagination of the host lipid membrane to form the parasitophorous vacuole; a regulated compartment to the parasite’s benefit that further shields it from the host-immune response (Caffaro and Boothroyd, 2011; Suarez et al., 2019). Lipid mediated signalling aids exocytosis of virulence factors from the rhoptries and micronemes, whereas both the host and parasite’s lipids are recruited to form the parasitophorous vacuole (Suarez et al., 2019; Katris et al., 2020). VLCFA play an important structural role in membrane biogenesis (Wang et al., 2018; Erdbrügger and Fröhlich, 2021) and several characterisation studies of the *T. gondii* lipidome have uncovered morphological aberrations during daughter cell formation and cytokinesis, incurred through loss of specific FA species (Amiar et al., 2016; Amiar et al., 2020; Dass et al., 2021; Onguka et al., 2021). Immunofluorescent microscopy and TEM found only a minor and not reproducible apicoplast morphology defect, and no effect on other organelles **(Supplementary figures 3 – 4)**, suggesting that the defect leading to the slower replication and reduced invasion does not seem to affect organelle morphology.

*Toxoplasma* bradyzoites are responsible for chronic infection and are resistant to drug treatment (Dunay Ildiko et al., 2018) which is why it is important to improve our understanding of their biology. Though their replication rate is slower than tachyzoites, *T. gondii* bradyzoites are not quiescent and must continue to scavenge resources from their hosts (Nolan et al., 2018; Kannan et al., 2021). We explored the contribution of FAE to early-stage differentiation *in vitro*, and brain cyst formation *in vivo*. Under alkaline media supplemented with 1% FBS, both ME49ΔELO-A and ME49ΔELO-AΔELO-B form cysts and successfully transition to the bradyzoite stage; marked by a depletion in *Tg*SAG1 expression and reciprocal increase in *Tg*BAG1 expression **(Figure 4G - H)**. Our study provides a snapshot of early-stage differentiation, and thus we cannot rule out an effect on spontaneous differentiation or in bradyzoite persistence.

Complementing previous *in vivo* CRISPR screen findings (Giuliano et al., 2023), we have shown in targeted deletions that *Tg*ELO enzymes are individually dispensable to *Toxoplasma in vitro* culture, but remain fitness conferring *in vivo*. Abrogation of *Tg*ELO-A and *Tg*ELO-B induced improvement of survival duration of infected BALB/c mice **(Figure 5D – E)**. Further, mice that survived till chronic infection onset had significantly reduced brain cyst burdens **(Figure 5F)**. *Tg*ELO enzymes are functionally homologous to those in *Saccharomyces cerevisiae* (Ramakrishnan et al., 2012), and depletion of corresponding ELO enzymes in *S. cerevisiae* induce reactive oxygen species sensitivity (Wang et al., 2018). It is possible that similar sensitivities result from *Tg*ELOs depletion and that the corresponding impact becomes relevant only when grown in the animal.

Our GC-MS bulk lipid analysis of tachyzoites was mostly concordant with the previously reported FAE interference studies. *Tg*ELO-A elongates C16:0 and C16:1 to C18:0 and C18:1, whereas *Tg*ELO-B elongates from C18:1 to C20:1, followed by C22:1 (Ramakrishnan et al., 2012; Ramakrishnan et al., 2015). Both ME49ΔELO-A and ME49ΔELO-AΔELO-B exhibited significant decreases in C18:0 and the cis isoform of C18:1, whereas C14:0, C16:0, and C16:1 significantly increased **(Figure 2A)**, indicative of a metabolic obstruction in their processing. Reflectively, C20:1 abundance was significantly lower and cis C18:1 was significantly higher in ME49ΔELO-AΔELO-B compared to ME49ΔELO-A **(Figure 2A)**. We explored the lipidome in our mutants grown under serum depleted conditions. Surprisingly, both *Tg*ELO deficient mutants demonstrated broadly similar ratios of relative LCFA abundance; the only exception being a significant increase in C18:0 in ME49ΔELO-AΔELO-B **(Figure 2B)**. Intriguingly, there was a small significant increase in trans C18:1 abundance in both mutants under 1% FBS conditions **(Figure 2B)** pointing towards increased FBS derived scavenging. Altogether, these experiments demonstrate that *T. gondii* remains viable under serum depletion despite impaired FAE and is capable of redirecting LCFA species to meet its requirements.

Glucose-derived carbon is used to generate LCFA precursors via *T. gondii’s* FASII and through labelling glucose in host-parasite co-cultures we can explore the incorporation of these carbons in LCFA. In principle, *T. gondii* can also scavenge FA synthesised from host-metabolised glucose; however, the majority of FA are likely to originate from the culture serum (Ramakrishnan et al., 2012; Grankvist et al., 2018), permitting comparative insights between parasite mutants. As expected, there was no significant difference in ^13^C labelling for the FASII products that fuel FAE, C14:0 and C16:0, between parental and mutants **(Figure 2C)**, demonstrating that apicoplast FASII remains functional when targeting ER-associated FAE. Likewise, *Tg*ELO-A and *Tg*ELO-B products, C18:1, and C20:1 demonstrated significantly reduced ^13^C labelling in both *Tg*ELO *T. gondii* mutants **(Figure 2C)**, indicative of expected reduced glycolytic carbon incorporation in these LCFA species. However, glycolytic carbon appeared to continue to integrate into other FAE products, where an unexpected significant increase in ^13^Clabelling occurred for multiple LCFA and VLCFA in either or both ME49ΔELO-A and ME49ΔELO-AΔELO-B **(Figure 2C)**.

Evidence that deletion of the *Tg*ELO units responsible for incorporating FA borne from glycolysis products, paradoxically increases glycolytic carbon incorporation in some LCFA has ramifications for our current understanding of FAE in *T. gondii*. LCFA species C18:1, C20:0, and C20:1 were broadly enriched in ME49 strain *T. gondii* than in ME49ΔELO-A and ME49ΔELO-AΔELO-B **(Supplementary figure 1F – H)**. Strikingly this was not the case for C22:0 and C24:0 whereby ME49ΔELO-AΔELO-B displayed higher levels of ^13^C incorporation **(Supplementary figure 1I)**. This finding can potentially be explained by three possible compensating factors: *Tg*ELO enzyme functional redundancy, increased parasite scavenging, or accessory pathways **(Figure 6)**.

*Tg*ELO-C was previously established to convert C22:1 to C26:1 with catalytic preference for host-derived over *de novo* synthesised unsaturated VLCFA (Ramakrishnan et al., 2012). Eukaryotic ELO-C homologues and paralogues commonly show functional overlap (Zheng et al., 2017; Wang et al., 2018; Pagura et al., 2023). Therefore, it is possible that *Tg*ELO-C could compensate ME49ΔELO-AΔELO-B through elongating scavenged FA and incorporating glycolytic carbons in FAE intermediates and products. Our phenotypic analyses demonstrating mutant sensitivity to serum availability *in vitro* **(Figure 3)** and increases in trans C18:1 abundance compared to ME49 *T. gondii* **(Figure 2A – B)**, supports this scenario.

**Figure 6:**
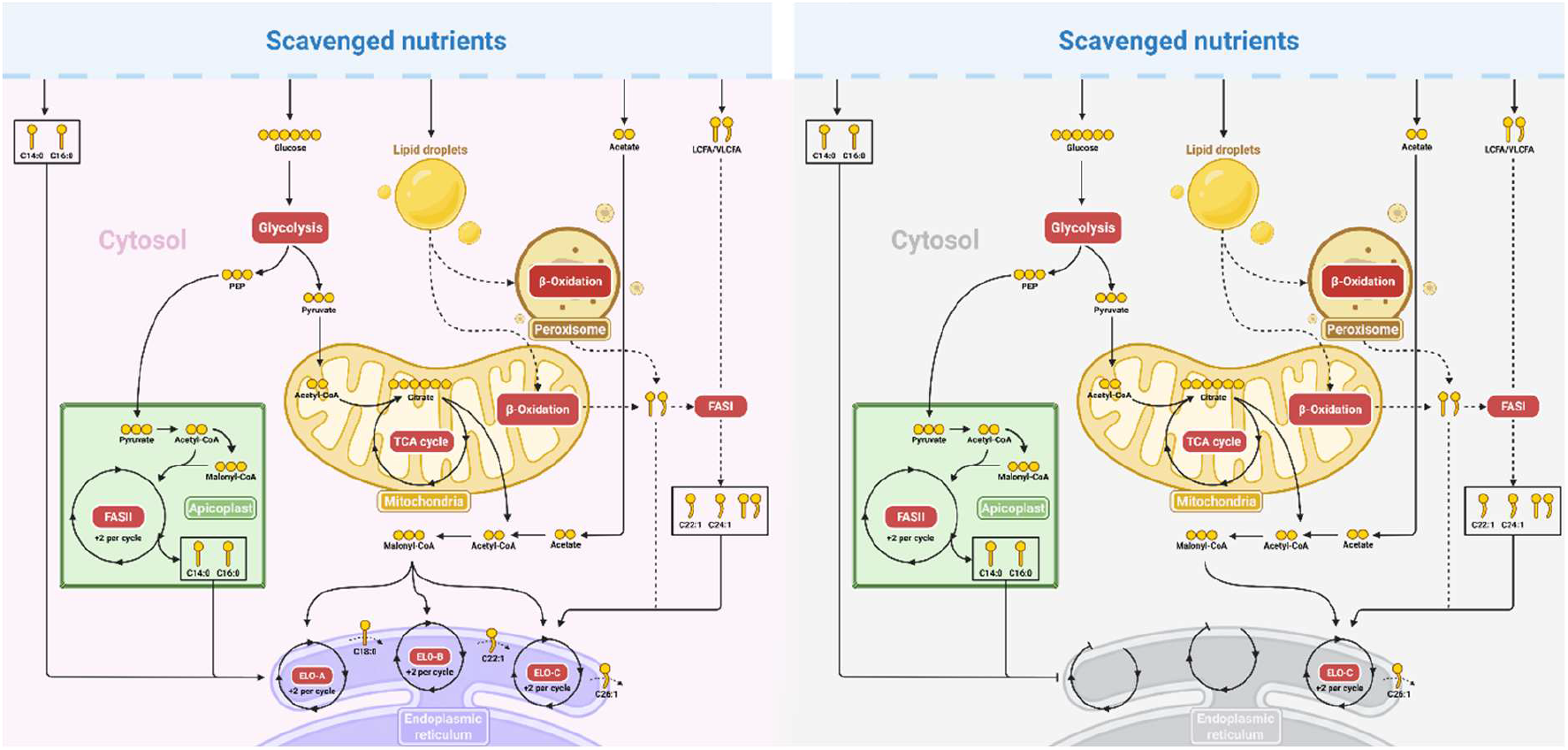
Proposed FAE pathway. A visual guide illustrating the pathways catabolised glucose funnels into FAE. Glucose derived carbon into LCFA and VLCFA production. Made with BioRender.com. Abbreviations: coenzyme A (CoA), fatty acid synthesis (FAS), Fatty Acid Elongation (FAE) long chain fatty acids (LCFA), tricarboxylic acid (TCA), very LCFA (VLCFA)

A gap in our understanding of lipid synthesis and metabolism in *T. gondii* exists with regards to extracellular lipid transport, the degradation of these lipids, and their subsequent mobilisation. *Tg*ACS3 has recently been implicated in funnelling acetyl-CoA into host derived LCFA C16:0 and C18:1 (Dass et al., 2024). A usual catabolitic pathway for FA is the FA β-oxidation, which typically occurs in mitochondria and peroxisomes, producing acetyl-CoA LCFA and VLCFA degradation (Ding et al., 2021). Despite *T. gondii* retaining enzymes indicative of this canonical pathway, its genome does not encode carnitine palmitoyltransferase 1, the canonical transporter responsible for shuttling of LCFA-derived acyl-CoA across the mitochondrial outer membrane (Nolan et al., 2018). Likewise, despite identification of peroxisome-associated FA β-oxidation protein orthologues, peroxisome-like structures have not observed in *T. gondii* (Moog et al., 2017; Charital et al., 2024b). However, recent work shows that the parasite possesses a bubblegum-like acytltransferase, TgACS1, which typically activates and transports FA to peroxisomal β-oxidation (Charital et al., 2024a). TgACS1 possesses a peroxisomal targeting sequence which targets the protein towards a specific location between the apicoplast and the Golgi apparatus that co-localises with catalase. The disruption of this enzyme affects parasite survival in low nutrient conditions, and most importantly, affects parasite gliding motility in extracellular stages under low nutrient content. This strongly suggest that the parasite could have an active β-oxidation pathway that is activatable under low nutrient content. Therefore, FA could be trafficked toward β-oxidation pathway for energetic reasons. The degradation products of these predicted pathways could be rerouted for increased glycolytic ^13^C incorporation in LCFA and VLCFA observed in *Tg*ELO deficient mutants.

Our findings may also be suggestive of greater modularity in *Toxoplasma* FAE to which other “opportunistic” enzymes, may contribute. Frameshift deletion of polyketide synthetase 1 (PKS1), or *Tg*FASI in RH strain *T. gondii* reportedly had no effect on parasite growth in DMEM cultures supplemented with 5% FCS, implying dispensability (Tymoshenko et al., 2015). *Tg*PKS2 is another FAS like complex but its catalytic function in *T. gondii* and other Apicomplexa is only partially resolved, and it remains to be seen if LCFA or VLCFA are its products or intermediates (Keeler et al., 2023; D’Ambrosio et al., 2023). Curiously, the mRNA expression of *Tg*PKS2 is upregulated in bradyzoites (D’Ambrosio et al., 2023) whereas *Tg*FASI mRNA expression is abundant in the unsporulated oocysts (Bushkin et al., 2013), and therefore these enzymes are speculated to play important roles in the lipid rich oocyst wall biogenesis (Bushkin et al., 2013; D’Ambrosio et al., 2023). Further research into the roles of these enzymes is required to explore their function in different life stages.

## Acknowledgments

The authors gratefully thank Ms Diane Vaughan, Mrs Alana Hamilton and Susan Baillie of the Cellular Analysis Facility at the University of Glasgow, for their support & assistance with the flow and light microscopy methods used in this work. The authors are grateful to Ms Maeve McLaughlin of the Queen Mary University of London Blizard Advanced Light Microscopy Facility for their advice and services. LS is supported by Wellcome Trust’s Wellcome Investigator Award (grant number 217173/Z/19/Z), and Wellcome Discovery Award (grant number 310879/Z/24/Z). CYB and YYB are supported by Agence Nationale de la Recherche, France (Project ApicoLipiAdapt grant ANR-21-CE44-0010; Project Apicolipidtraffic grant ANR-23-CE15-0009-01; Project OIL grant ANR-24-CE15-2171-02; Project Plasmohost ANR-25-CE11-3606), The Fondation pour la Recherche Médicale (FRM EQU202103012700), Laboratoire d’Excellence Parafrap, France (grant ANR-11-LABX-0024), LIA-IRP CNRS Program (Apicolipid project), the Université Grenoble Alpes (IDEX ISP Apicolipid) and Région Auvergne Rhone-Alpes for the lipidomics analyses platform (Grant IRICE Project GEMELI), Collaborative Research Program Grant CEFIPRA (Project 6003-1), IARDP grant (#2023-0414) by the CEFIPRA (MESRI-DBT). DS is funded by a Moredun Research Fellowship from The Moredun Foundation and through the Scottish Government Rural and Environmental Science and Analytical Services Division (RESAS) strategic research programme. SC is supported by an Industrial Partnership PhD programme studentship between University of Glasgow College of Medical, Veterinary and Life Sciences and Moredun Research Institute.

## Supplementary materials

**Table 1:**
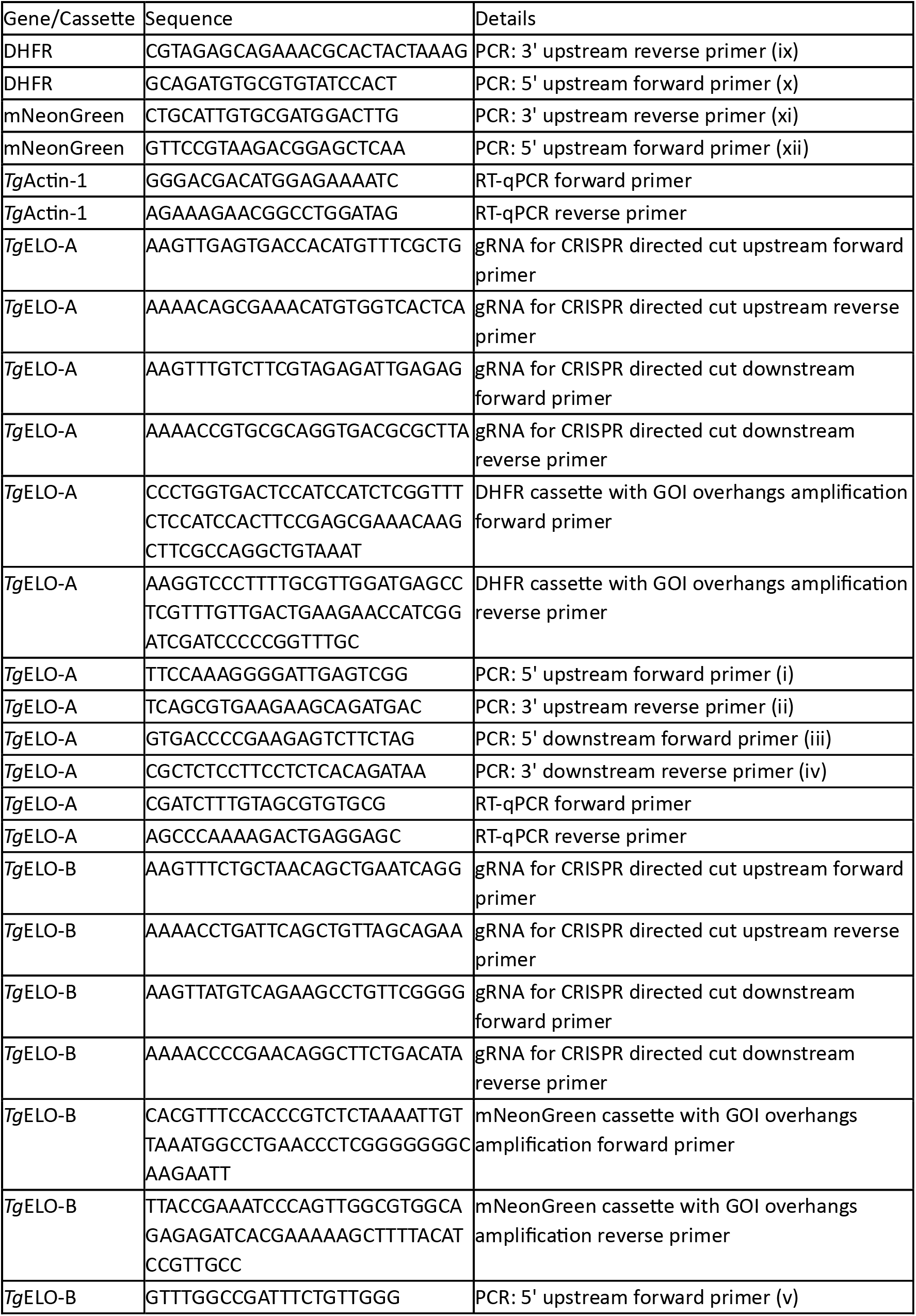
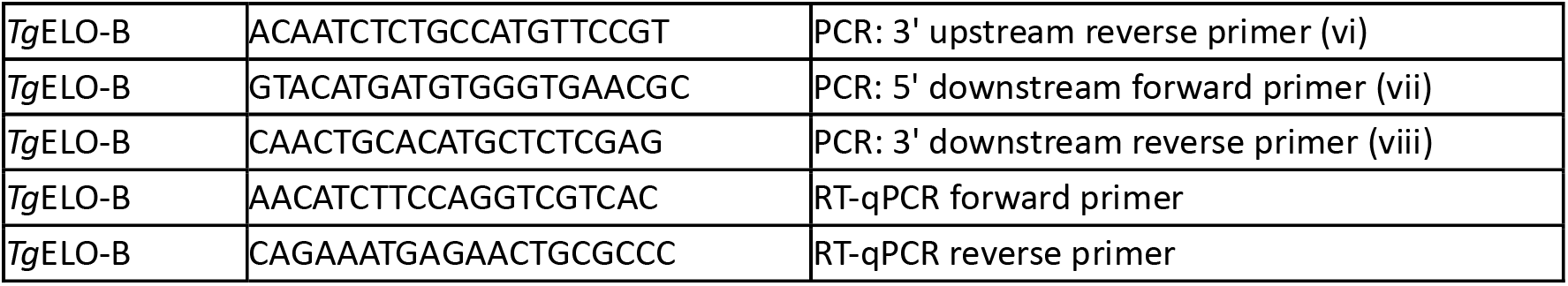
List of oligonucleotides used in the generation and validation of ME49ΔELO-A and ME49ΔELO-AΔELO-B.

**Supplementary figure 1:**
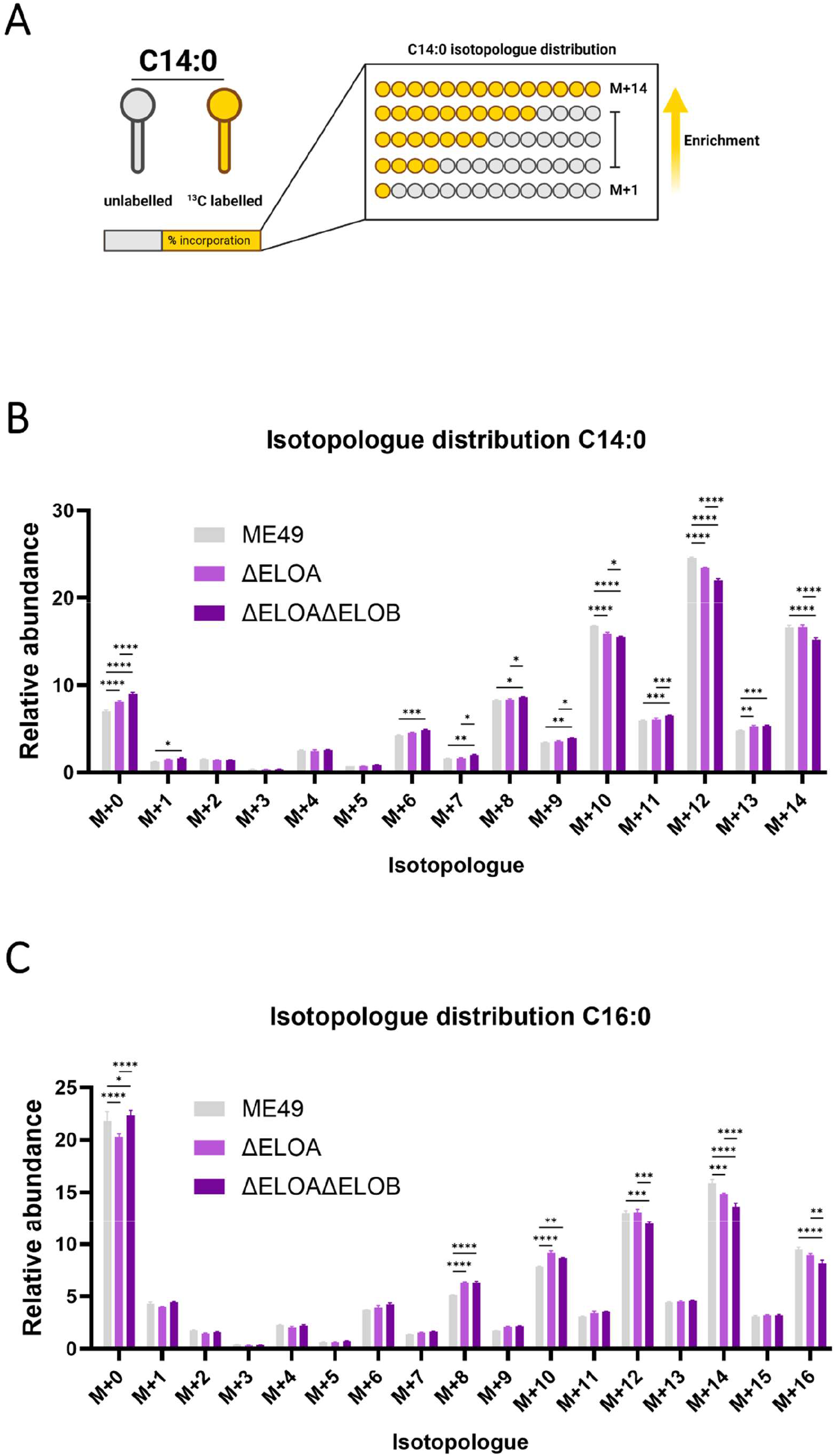
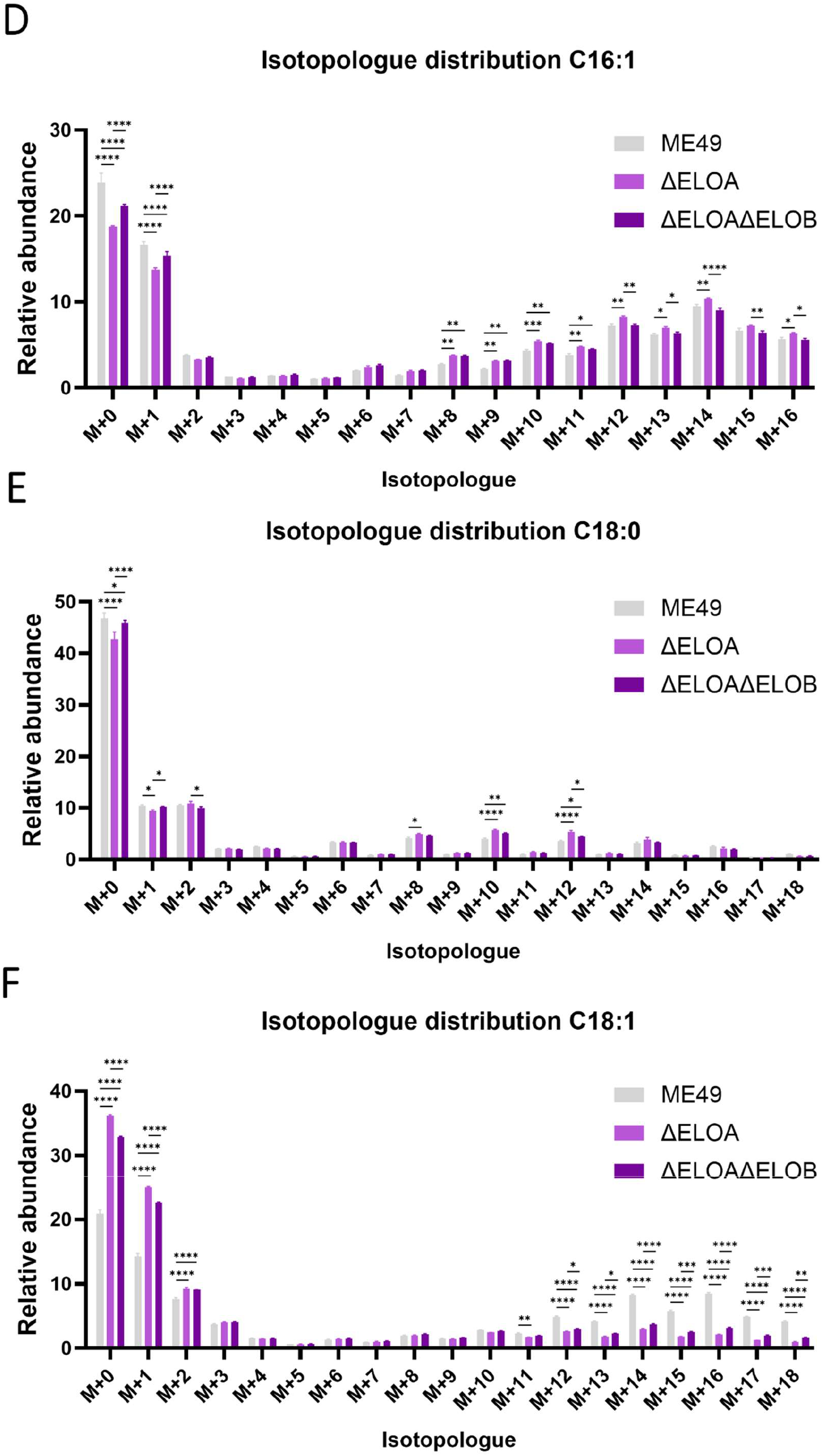
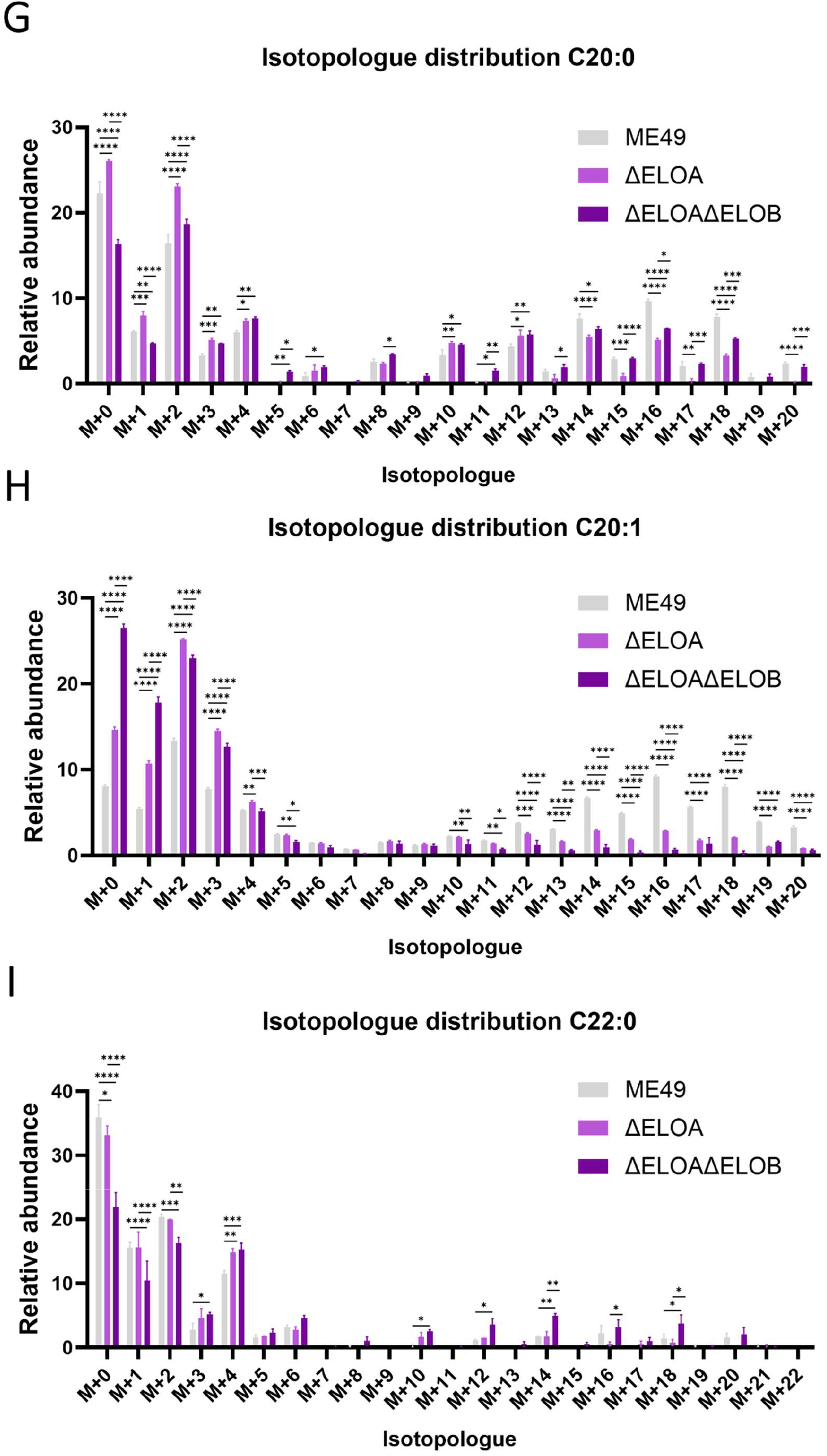
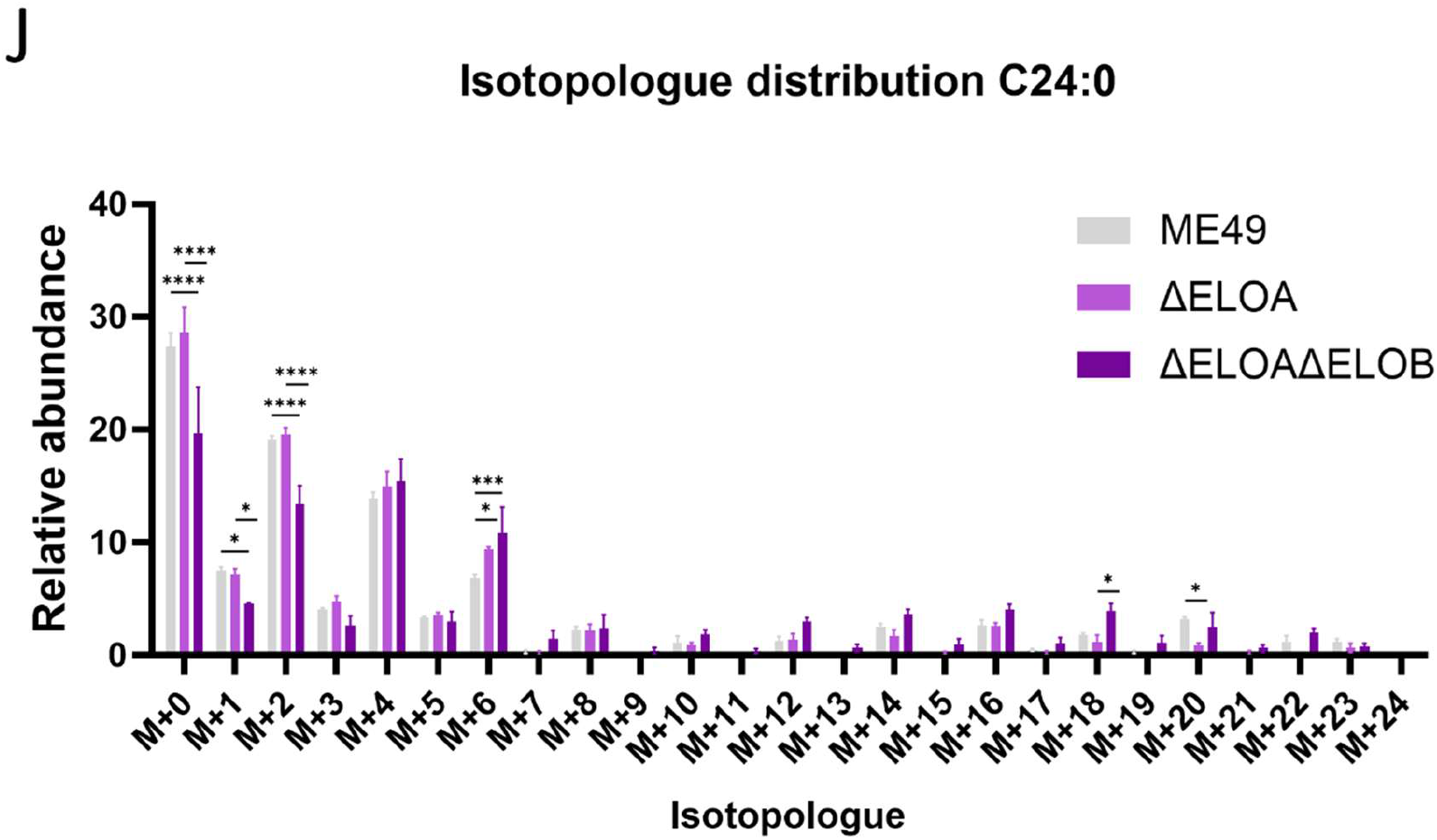
**(A)** Visual aid for metabolite labelling experiment results interpretation using C14:0 as an example. Mass isotopologue distribution of **(B)** C14:0, **(C)** C16:0, **(D)** C16:1, **(E)** C18:0, **(F)** C18:1, **(G)** C20:0, and **(H)** C20:1 LCFA species, as well as **(I)** C22:0 and **(J)** C24:0 VLCFA species in ME49, ME49ΔELO-A, and ME49ΔELO-AΔELO-B *T. gondii*. Intracellular and extracellular parasites were quenched at 0 °C, prior to processing for GC-MS. Analysis via Two-way ANOVA with Fishers LSD, n = 3. Unpaired t-test values: * P < 0.05, ** P < 0.01, *** P < 0.001, **** P < 0.0001.

**Supplementary figure 2:**
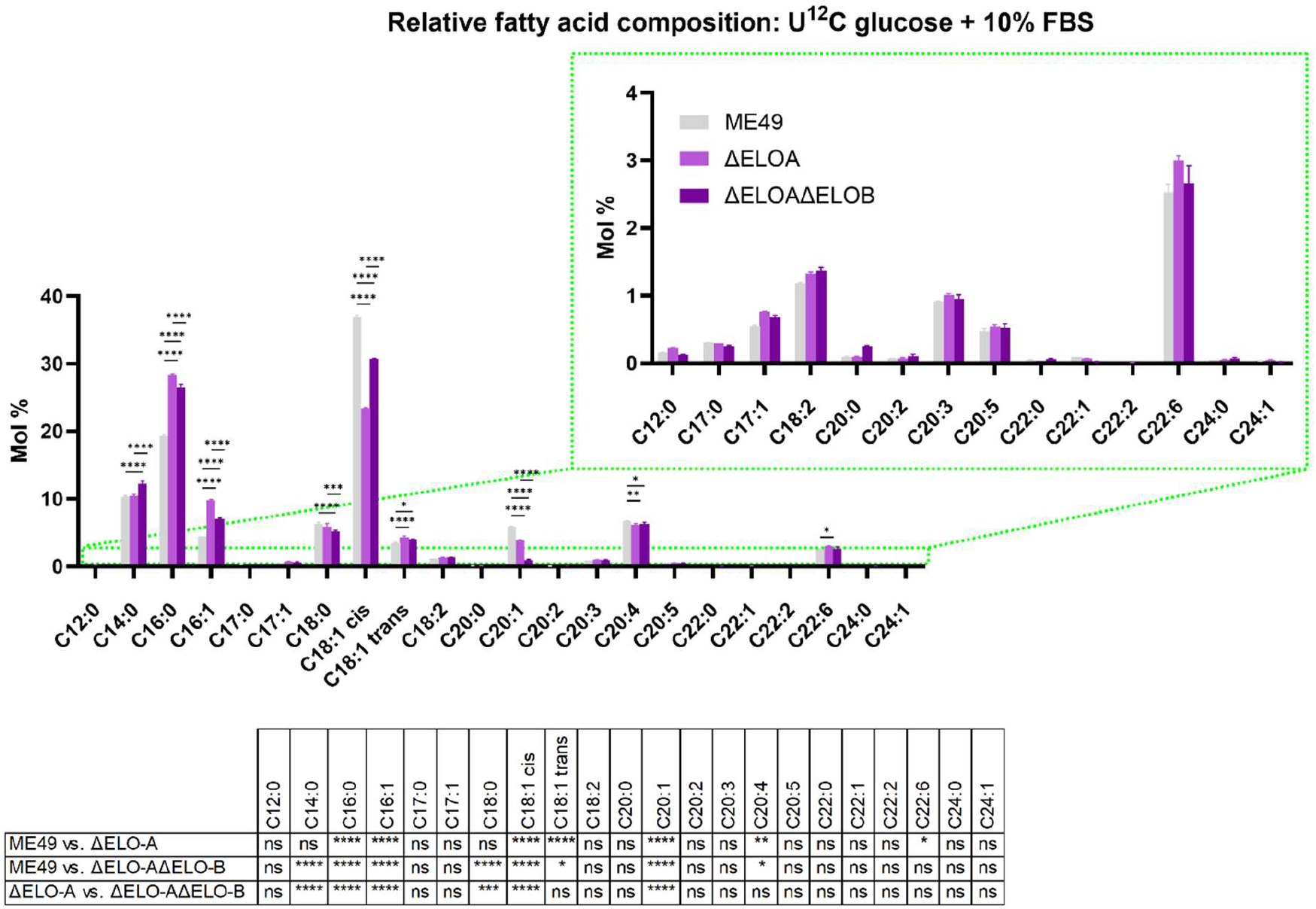
Relative FA composition in ME49 strain T. gondii and derived TgELO depleted mutants in percentage molarity (Mol %). Parasites were cultured in DMEM (11966) supplemented with U12C-glucose, with 10% FBS, and harvested when confluent. Intracellular and extracellular parasites were quenched at 0 °C, prior to processing for gas chromatography-mass spectroscopy (GC-MS). Analysis via Two-way ANOVA with Fishers LSD, n = 3. Unpaired t-test values: * P < 0.05, ** P < 0.01, *** P < 0.001, **** P < 0.0001, (ns) not significant.

**Supplementary figure 3:**
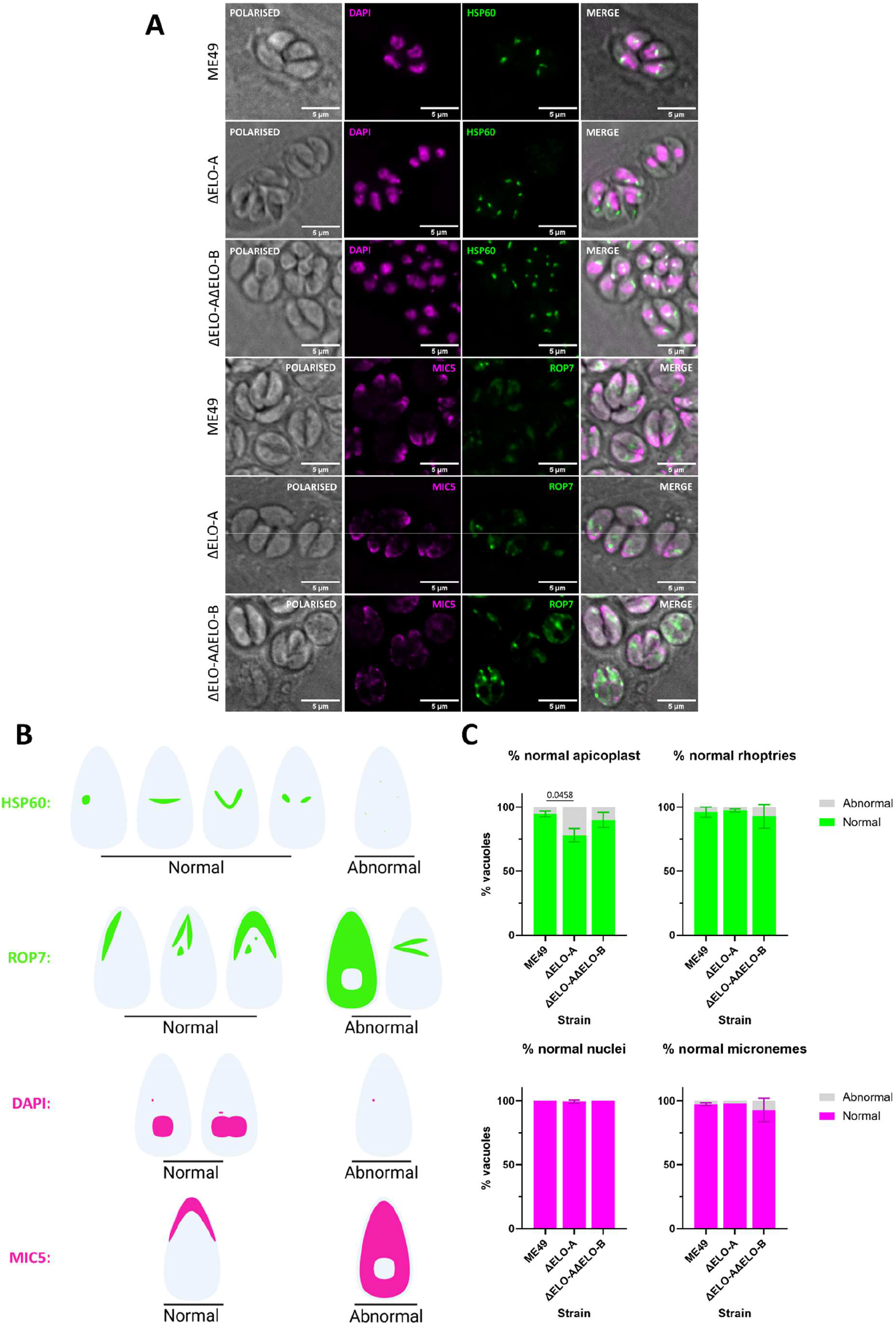
**(A)** Immunofluorescent staining of *T. gondii* organelles: nucleus, apicoplast, micronemes, rhoptries. Targets were selected on perceived importance to parasite replication and invasion. ME49, ME49ΔELO-A, and ME49ΔELO-AΔELO-B were co-cultured with host HFF-1 in DMEM (11965) supplemented with 1% FBS for 36 hrs prior to fixation. Genomic material and apicoplast were stained with DAPI and anti-HSP60, whereas micronemes and rhoptries were stained with anti-MIC5 and anti-ROP7 respectively. Images shown are Z-projections where brightness, contrast settings were normalised to a representative acquired image of ME49 *T. gondii*. **(B)** Organelle morphs were categorised as normal or abnormal based on literature descriptions and **(C)** percentage of vacuoles categorised as such were quantified. Analysis via Kruskal Wallace one-way ANOVA with Dunn’s post hoc. Error bars are Mean ± SD, representative of three biological replicates, where n=50 vacuoles were assessed.

**Supplementary figure 4:**
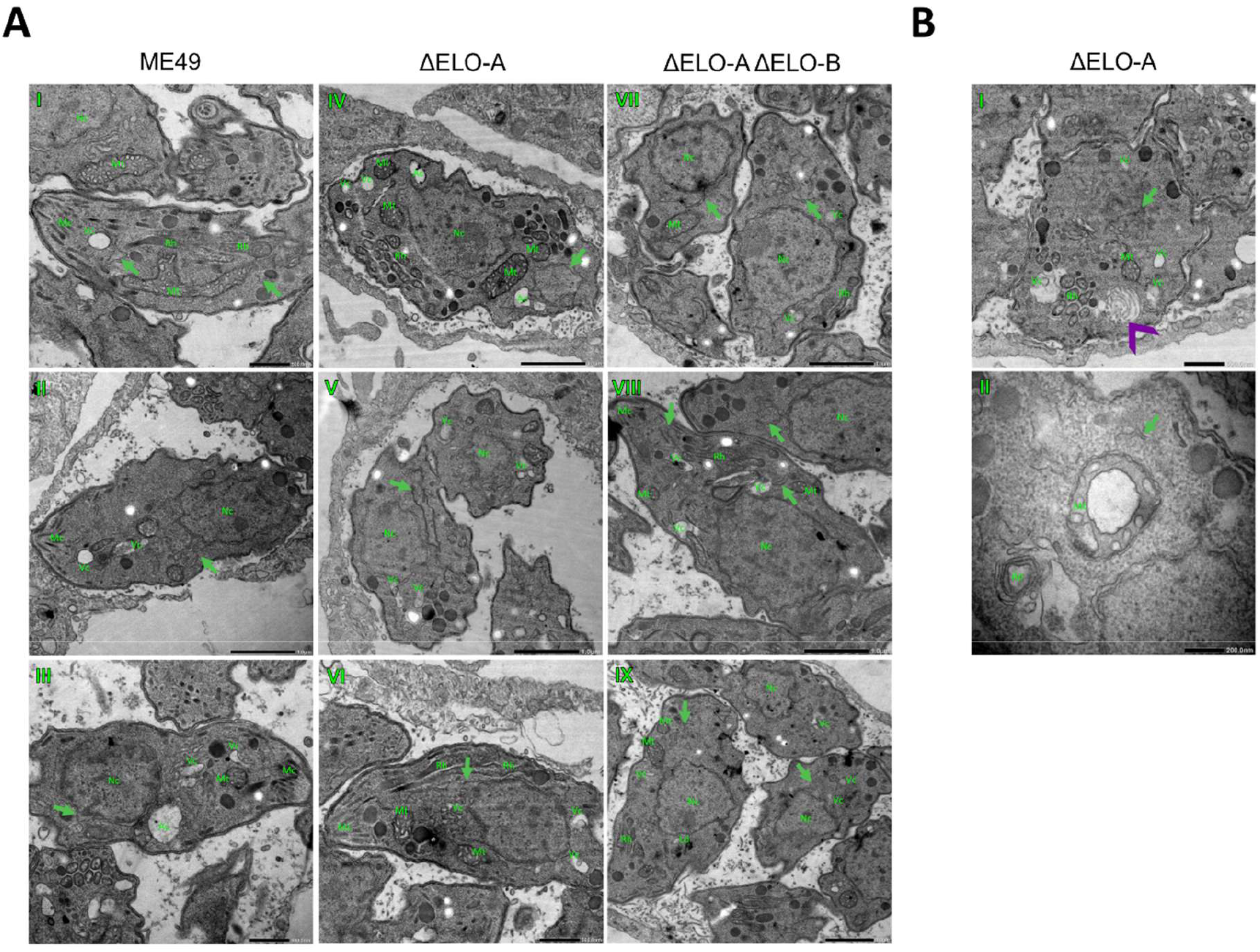
**(A)** Representative electron microscopy images of intracellular (AI – III) ME49 T. gondii, (AIV – VI) ME49ΔELO-A, and (AVII – IX) ME49Δ ELO-AΔELO-B. **(B)** Anomalous phenotypes observed in ME49ΔELO-A including (BI) a multilamellar structure and (BII) a compartment within a mitochondrion. Host HFF-1 and *T. gondii* cells were co-cultured in DMEM (11965) supplemented with 1% FBS for three days before being fixed and processed for imaging. Scale bars shown ranges from 1000 – 200nm. Green arrows indicate Endoplasmic Reticulum. Purple chevron indicates multilamellar structure. Abbreviations: acidocalcisome (Ac), micronemes (Mc), mitochondria (Mt), nucleus (Nc), rhoptries (Rh), vacuole (Vc).

## References

Aghabi, D., Sloan, M., Gill, G., Hartmann, E., Antipova, O., Dou, Z., Guerra, A. J., Carruthers, V. B. & Harding, C. R. 2023. The vacuolar iron transporter mediates iron detoxification in Toxoplasma gondii. Nature Communications, 14(1), pp 3659.

Alvarez-Jarreta, J., Amos, B., Aurrecoechea, C., Bah, S., Barba, M., Barreto, A., Basenko, E. Y., Belnap, R., Blevins, A., Böhme, U., Brestelli, J., Brown, S., Callan, D., Campbell, L. I., Christophides, G. K., Crouch, K., Davison, H. R., DeBarry, J. D., Demko, R., Doherty, R., Duan, Y., Dundore, W., Dyer, S., Falke, D., Fischer, S., Gajria, B., Galdi, D., Giraldo-Calderón, G. I., Harb, O. S., Harper, E., Helb, D., Howington, C., Hu, S., Humphrey, J., Iodice, J., Jones, A., Judkins, J., Kelly, S. A., Kissinger, J. C., Kittur, N., Kwon, D. K., Lamoureux, K., Li, W., Lodha, D., MacCallum, R. M., Maslen, G., McDowell, M. A., Myers, J., Nural, M. V., Roos, D. S., Rund, S. S. C., Shanmugasundram, A., Sitnik, V., Spruill, D., Starns, D., Tomko, S. S., Wang, H., Warrenfeltz, S., Wieck, R., Wilkinson, P. A. & Zheng, J. 2024. VEuPathDB: the eukaryotic pathogen, vector and host bioinformatics resource center in 2023. Nucleic Acids Research, 52(D1), pp D808–D816.

Amiar, S., Katris, N. J., Berry, L., Dass, S., Duley, S., Arnold, C.-S., Shears, M. J., Brunet, C., Touquet, B., McFadden, G. I., Yamaryo-Botté, Y. & Botté, C. Y. 2020. Division and Adaptation to Host Environment of Apicomplexan Parasites Depend on Apicoplast Lipid Metabolic Plasticity and Host Organelle Remodeling. Cell Reports, 30(11), pp 3778–3792.e9.

Amiar, S., MacRae, J. I., Callahan, D. L., Dubois, D., van Dooren, G. G., Shears, M. J., Cesbron-Delauw, M.-F., Maréchal, E., McConville, M. J., McFadden, G. I., Yamaryo-Botté, Y. & Botté, C. Y. 2016. Apicoplast-Localized Lysophosphatidic Acid Precursor Assembly Is Required for Bulk Phospholipid Synthesis in Toxoplasma gondii and Relies on an Algal/Plant-Like Glycerol 3-Phosphate Acyltransferase. PLOS Pathogens, 12(8), pp e1005765.

Arnold, C.-S., Alazzi, A.-M., Shunmugam, S., Janouškovec, J., Berry, L., Charital, S., Gautier, T., Duley, S., Jublot, D., Lemaire-Vieille, C., Cesbron-Delauw, M.-F., Cavailles, P., Govin, J., Katris, N. J., Yamaryo-Botté, Y. & Botté, C. Y. 2025. A P5-ATPase, TgFLP12, diverging from plant chloroplast lipid transporters mediates apicoplast fatty export in Toxoplasma. Nature Communications, 16(1), pp 5538.

Ben Chaabene, R., Martinez, M., Bonavoglia, A., Maco, B., Chang, Y. W., Lentini, G. & Soldati-Favre, D. 2024. Toxoplasma gondii rhoptry discharge factor 3 is essential for invasion and microtubule-associated vesicle biogenesis. PLoS Biol, 22(8), pp e3002745.

Bitew, M. A., Paredes-Santos, T. C., Maru, P., Krishnamurthy, S., Wang, Y., Sangaré, L. O., Duley, S., Yamaryo-Botté, Y., Botté, C. Y. & Saeij, J. P. J. 2025. A genome-wide CRISPR screen identifies GRA38 as a key regulator of lipid homeostasis during Toxoplasma gondii adaptation to lipid-rich conditions. Nature Communications, 16(1), pp 11177.

Blader, I. J., Coleman, B. I., Chen, C. T. & Gubbels, M. J. 2015. Lytic Cycle of Toxoplasma gondii: 15 Years Later. Annu Rev Microbiol, 69(463–85.

Bohne, W., Gross, U., Ferguson, D. J. P. & Heesemann, J. 1995. Cloning and characterization of a bradyzoite-specifically expressed gene (hsp30/bag1) of Toxoplasma gondii, related to genes encoding small heat-shock proteins of plants. Molecular Microbiology, 16(6), pp 1221–1230.

Botté, C. Y., Yamaryo-Botté, Y., Rupasinghe, T. W. T., Mullin, K. A., MacRae, J. I., Spurck, T. P., Kalanon, M., Shears, M. J., Coppel, R. L., Crellin, P. K., Maréchal, E., McConville, M. J. & McFadden, G. I. 2013. Atypical lipid composition in the purified relict plastid (apicoplast) of malaria parasites. Proceedings of the National Academy of Sciences, 110(18), pp 7506.

Breinich, M. S., Ferguson, D. J. P., Foth, B. J., van Dooren, G. G., Lebrun, M., Quon, D. V., Striepen, B., Bradley, P. J., Frischknecht, F., Carruthers, V. B. & Meissner, M. 2009. A Dynamin Is Required for the Biogenesis of Secretory Organelles in Toxoplasma gondii. Current Biology, 19(4), pp 277–286.

Bushkin, G. G., Motari, E., Carpentieri, A., Dubey Jitender, P., Costello Catherine, E., Robbins Phillips, W. & Samuelson, J. 2013. Evidence for a Structural Role for Acid-Fast Lipids in Oocyst Walls of Cryptosporidium, Toxoplasma, and Eimeria. mBio, 4(5), pp 10.1128/mbio.00387-13.

Caffaro, C. E. & Boothroyd, J. C. 2011. Evidence for host cells as the major contributor of lipids in the intravacuolar network of Toxoplasma-infected cells. Eukaryot Cell, 10(8), pp 1095–9.

Charital, S., Lourdel, A., Quansah, N., Botté, C. Y. & Yamaryo-Botté, Y. 2024a. Monitoring of Lipid Fluxes Between Host and Plastid-Bearing Apicomplexan Parasites. Methods Mol Biol, 2776(197-204.

Charital, S., Shunmugam, S., Dass, S., Alazzi Anna, M., Arnold, C.-S., Katris Nicholas, J., Duley, S., Quansah Nyamekye, A., Pierrel, F., Govin, J., Yamaryo-Botté, Y. & Botté Cyrille, Y. 2024b. The acyl-CoA synthetase TgACS1 allows neutral lipid metabolism and extracellular motility in Toxoplasma gondii through relocation via its peroxisomal targeting sequence (PTS) under low nutrient conditions. mBio, 15(4), pp e00427–24.

Chen, K., Günay-Esiyok, Ö., Klingeberg, M., Marquardt, S., Pomorski, T. G. & Gupta, N. 2021. Aminoglycerophospholipid flipping and P4-ATPases in Toxoplasma gondii. Journal of Biological Chemistry, 296(100315.

Chen, M., Koszti, S. G., Bonavoglia, A., Maco, B., von Rohr, O., Peng, H.-J., Soldati-Favre, D. & Kloehn, J. 2025. Dissecting apicoplast functions through continuous cultivation of Toxoplasma gondii devoid of the organelle. Nature Communications, 16(1), pp 2095.

Curt-Varesano, A., Braun, L., Ranquet, C., Hakimi, M.-A. & Bougdour, A. 2016. The aspartyl protease TgASP5 mediates the export of the Toxoplasma GRA16 and GRA24 effectors into host cells. Cellular Microbiology, 18(2), pp 151–167.

D’Ambrosio, H. K., Keeler, A. M. & Derbyshire, E. R. 2023. Examination of Secondary Metabolite Biosynthesis in Apicomplexa. ChemBioChem, 24(17), pp e202300263.

Dass, S., Shunmugam, S., Berry, L., Arnold, C.-S., Katris, N. J., Duley, S., Pierrel, F., Cesbron-Delauw, M.-F., Yamaryo-Botté, Y. & Botté, C. Y. 2021. Toxoplasma LIPIN is essential in channeling host lipid fluxes through membrane biogenesis and lipid storage. Nature Communications, 12(1), pp 2813.

Dass, S., Shunmugam, S., Charital, S., Duley, S., Arnold, C.-S., Katris, N. J., Cavaillès, P., Cesbron-Delauw, M.-F., Yamaryo-Botté, Y. & Botté, C. Y. 2024. Toxoplasma acyl-CoA synthetase *Tg*ACS3 is crucial to channel host fatty acids in lipid droplets and for parasite propagation. Journal of Lipid Research, 65(10), pp.

DeRocher, A. E., Coppens, I., Karnataki, A., Gilbert, L. A., Rome, M. E., Feagin, J. E., Bradley, P. J. & Parsons, M. 2008. A thioredoxin family protein of the apicoplast periphery identifies abundant candidate transport vesicles in Toxoplasma gondii. Eukaryot Cell, 7(9), pp 1518–29.

Di Cristina, M., Dou, Z., Lunghi, M., Kannan, G., Huynh, M.-H., McGovern, O. L., Schultz, T. L., Schultz, A. J., Miller, A. J., Hayes, B. M., van der Linden, W., Emiliani, C., Bogyo, M., Besteiro, S., Coppens, I. & Carruthers, V. B. 2017. Toxoplasma depends on lysosomal consumption of autophagosomes for persistent infection. Nature Microbiology, 2(8), pp 17096.

Ding, L., Sun, W., Balaz, M., He, A., Klug, M., Wieland, S., Caiazzo, R., Raverdy, V., Pattou, F., Lefebvre, P., Lodhi, I. J., Staels, B., Heim, M. & Wolfrum, C. 2021. Peroxisomal β-oxidation acts as a sensor for intracellular fatty acids and regulates lipolysis. Nature Metabolism, 3(12), pp 1648–1661.

Dogga, S. K., Mukherjee, B., Jacot, D., Kockmann, T., Molino, L., Hammoudi, P.-M., Hartkoorn, R. C., Hehl, A. B. & Soldati-Favre, D. 2017. A druggable secretory protein maturase of Toxoplasma essential for invasion and egress. eLife, 6(e27480.

Dubois, D., Fernandes, S., Amiar, S., Dass, S., Katris, N. J., Botté, C. Y. & Yamaryo-Botté, Y. 2018. Toxoplasma gondii acetyl-CoA synthetase is involved in fatty acid elongation (of long fatty acid chains) during tachyzoite life stages. Journal of Lipid Research, 59(6), pp 994–1004.

Dunay Ildiko, R., Gajurel, K., Dhakal, R., Liesenfeld, O. & Montoya Jose, G. 2018. Treatment of Toxoplasmosis: Historical Perspective, Animal Models, and Current Clinical Practice. Clinical Microbiology Reviews, 31(4), pp 10.1128/cmr.00057-17.

Erdbrügger, P. & Fröhlich, F. 2021. The role of very long chain fatty acids in yeast physiology and human diseases. 402(1), pp 25–38.

Fernández-Escobar, M., Schares, G., Maksimov, P., Joeres, M., Ortega-Mora, L. M. & Calero-Bernal, R. 2022. Toxoplasma gondii Genotyping: A Closer Look Into Europe. Frontiers in Cellular and Infection Microbiology, Volume 12 - 2022(

Giuliano, C. J., Wei, K. J., Harling, F. M., Waldman, B. S., Farringer, M. A., Boydston, E. A., Lan, T. C. T., Thomas, R. W., Herneisen, A. L., Sanderlin, A. G., Coppens, I., Dvorin, J. D. & Lourido, S. 2023. Functional profiling of the Toxoplasma genome during acute mouse infection. bioRxiv.

Grankvist, N., Watrous, J. D., Lagerborg, K. A., Lyutvinskiy, Y., Jain, M. & Nilsson, R. 2018. Profiling the Metabolism of Human Cells by Deep 13C Labeling. Cell Chemical Biology, 25(11), pp 1419–1427.e4.

Hajimohammadi, B., Ahmadian, S., Firoozi, Z., Askari, M., Mohammadi, M., Eslami, G., Askari, V., Loni, E., Barzegar-Bafrouei, R. & Boozhmehrani, M. J. 2022. A Meta-Analysis of the Prevalence of Toxoplasmosis in Livestock and Poultry Worldwide. EcoHealth, 19(1), pp 55–74.

Hosseini, S. A., Amouei, A., Sharif, M., Sarvi, S., Galal, L., Javidnia, J., Pagheh, A. S., Gholami, S., Mizani, A. & Daryani, A. 2018. Human toxoplasmosis: a systematic review for genetic diversity of Toxoplasma gondii in clinical samples. Epidemiol Infect, 147(e36.

Jacot, D., Daher, W. & Soldati-Favre, D. 2013. Toxoplasma gondii myosin F, an essential motor for centrosomes positioning and apicoplast inheritance. The EMBO Journal, 32(12), pp 1702–1716.

Kannan, G., Thaprawat, P., Schultz Tracey, L. & Carruthers Vern, B. 2021. Acquisition of Host Cytosolic Protein by Toxoplasma gondii Bradyzoites. mSphere, 6(1), pp 10.1128/msphere.00934-20.

Katris, N. J., Yamaryo-Botte, Y., Janouškovec, J., Shunmugam, S., Arnold, C.-S., Yang, A. S. P., Vardakis, A., Stewart, R. J., Sauerwein, R., McFadden, G. I., Tonkin, C. J., Cesbron-Delauw, M.-F., Waller, R. F. & Botte, C. Y. 2020. Rapid kinetics of lipid second messengers controlled by a cGMP signalling network coordinates apical complex functions in Toxoplasma tachyzoites. bioRxiv, 2020.06.19.160341.

Keeler, A. M., D’Ambrosio, H. K., Ganley, J. G. & Derbyshire, E. R. 2023. Characterization of Unexpected Self-Acylation Activity of Acyl Carrier Proteins in a Modular Type I Apicomplexan Polyketide Synthase. ACS Chemical Biology, 18(4), pp 785–793.

Kim, K. & Weiss, L. M. 2004. Toxoplasma gondii: the model apicomplexan. International Journal for Parasitology, 34(3), pp 423–432.

Kloehn, J., Lunghi, M., Varesio, E., Dubois, D. & Soldati-Favre, D. 2021. Untargeted Metabolomics Uncovers the Essential Lysine Transporter in Toxoplasma gondii. Metabolites, 11(8), pp.

Kloehn, J., Oppenheim, R. D., Siddiqui, G., De Bock, P.-J., Kumar Dogga, S., Coute, Y., Hakimi, M.-A., Creek, D. J. & Soldati-Favre, D. 2020. Multi-omics analysis delineates the distinct functions of sub-cellular acetyl-CoA pools in Toxoplasma gondii. BMC Biology, 18(1), pp 67.

Kremer, K., Kamin, D., Rittweger, E., Wilkes, J., Flammer, H., Mahler, S., Heng, J., Tonkin, C. J., Langsley, G., Hell, S. W., Carruthers, V. B., Ferguson, D. J. P. & Meissner, M. 2013. An Overexpression Screen of Toxoplasma gondii Rab-GTPases Reveals Distinct Transport Routes to the Micronemes. PLOS Pathogens, 9(3), pp e1003213.

Krishnan, A., Kloehn, J., Lunghi, M., Chiappino-Pepe, A., Waldman, B. S., Nicolas, D., Varesio, E., Hehl, A., Lourido, S., Hatzimanikatis, V. & Soldati-Favre, D. 2020. Functional and Computational Genomics Reveal Unprecedented Flexibility in Stage-Specific Toxoplasma Metabolism. Cell Host & Microbe, 27(2), pp 290–306.e11.

Labun, K., Montague, T. G., Krause, M., Torres Cleuren, Y. N., Tjeldnes, H. & Valen, E. 2019. CHOPCHOP v3: expanding the CRISPR web toolbox beyond genome editing. Nucleic Acids Research, 47(W1), pp W171–W174.

Lévêque, M. F., Berry, L., Yamaryo-Botté, Y., Nguyen, H. M., Galera, M., Botté, C. Y. & Besteiro, S. 2017. TgPL2, a patatin-like phospholipase domain-containing protein, is involved in the maintenance of apicoplast lipids homeostasis in Toxoplasma. Molecular Microbiology, 105(1), pp 158–174.

Li, Y., Liu, Y., Xiu, F., Wang, J., Cong, H., He, S., Shi, Y., Wang, X., Li, X. & Zhou, H. 2018. Characterization of exosomes derived from Toxoplasma gondii and their functions in modulating immune responses. International Journal of Nanomedicine, 13(null), pp 467–477.

Liang, X., Cui, J., Yang, X., Xia, N., Li, Y., Zhao, J., Gupta, N. & Shen, B. 2020. Acquisition of exogenous fatty acids renders apicoplast-based biosynthesis dispensable in tachyzoites of *Toxoplasma*. Journal of Biological Chemistry, 295(22), pp 7743–7752.

Lim, L., Linka, M., Mullin, K. A., Weber, A. P. M. & McFadden, G. I. 2010. The carbon and energy sources of the non-photosynthetic plastid in the malaria parasite. FEBS Letters, 584(3), pp 549–554.

Livak, K. J. & Schmittgen, T. D. 2001. Analysis of Relative Gene Expression Data Using Real-Time Quantitative PCR and the 2−ΔΔCT Method. Methods, 25(4), pp 402–408.

Manger, I., D., Hehl, A., B. & Boothroyd, J., C. 1998. The Surface of Toxoplasma Tachyzoites Is Dominated by a Family of Glycosylphosphatidylinositol-Anchored Antigens Related to SAG1. Infection and Immunity, 66(5), pp 2237–2244.

Mazumdar, J., H. Wilson, E., Masek, K., A. Hunter, C. & Striepen, B. 2006. Apicoplast fatty acid synthesis is essential for organelle biogenesis and parasite survival in Toxoplasma gondii. Proceedings of the National Academy of Sciences, 103(35), pp 13192–13197.

Molan, A., Nosaka, K., Hunter, M. & Wang, W. 2019. Global status of Toxoplasma gondii infection: systematic review and prevalence snapshots. Trop Biomed, 36(4), pp 898–925.

Montazeri, M., Mikaeili Galeh, T., Moosazadeh, M., Sarvi, S., Dodangeh, S., Javidnia, J., Sharif, M. & Daryani, A. 2020. The global serological prevalence of Toxoplasma gondii in felids during the last five decades (1967–2017): a systematic review and meta-analysis. Parasites & Vectors, 13(1), pp 82.

Moog, D., Przyborski, J. M. & Maier, U. G. 2017. Genomic and Proteomic Evidence for the Presence of a Peroxisome in the Apicomplexan Parasite Toxoplasma gondii and Other Coccidia. Genome Biology and Evolution, 9(11), pp 3108–3121.

Morlon-Guyot, J., Berry, L., Sauquet, I., Singh Pall, G., El Hajj, H., Meissner, M. & Daher, W. 2018. Conditional knock-down of a novel coccidian protein leads to the formation of aberrant apical organelles and abrogates mature rhoptry positioning in Toxoplasma gondii. Molecular and Biochemical Parasitology, 223(19–30.

Mullin, K. A., Lim, L., Ralph, S. A., Spurck, T. P., Handman, E. & McFadden, G. I. 2006. Membrane transporters in the relict plastid of malaria parasites. Proc Natl Acad Sci U S A, 103(25), pp 9572–7.

Nolan, S. J., Romano, J. D., Kline, J. T. & Coppens, I. 2018. Novel Approaches To Kill Toxoplasma gondii by Exploiting the Uncontrolled Uptake of Unsaturated Fatty Acids and Vulnerability to Lipid Storage Inhibition of the Parasite. Antimicrob Agents Chemother, 62(10), pp.

Onguka, O., Babin, B. M., Lakemeyer, M., Foe, I. T., Amara, N., Terrell, S. M., Lum, K. M., Cieplak, P., Niphakis, M. J., Long, J. Z. & Bogyo, M. 2021. Toxoplasma gondii serine hydrolases regulate parasite lipid mobilization during growth and replication within the host. Cell Chemical Biology, 28(10), pp 1501–1513.e5.

Oppenheim, R. D., Creek, D. J., Macrae, J. I., Modrzynska, K. K., Pino, P., Limenitakis, J., Polonais, V., Seeber, F., Barrett, M. P., Billker, O., McConville, M. J. & Soldati-Favre, D. 2014. BCKDH: The Missing Link in Apicomplexan Mitochondrial Metabolism Is Required for Full Virulence of Toxoplasma gondii and Plasmodium berghei. PLOS Pathogens, 10(7), pp e1004263.

Pagura, L., Dumoulin, P. C., Ellis, C. C., Mendes, M. T., Estevao, I. L., Almeida, I. C. & Burleigh, B. A. 2023. Fatty acid elongases 1-3 have distinct roles in mitochondrial function, growth, and lipid homeostasis in Trypanosoma cruzi. Journal of Biological Chemistry, 299(6), pp 104715.

Pasquarelli, R. R., Quan, J. J., Cheng, E. S., Yang, V., Britton, T. A., Sha, J., Wohlschlegel, J. A. & Bradley, P. J. 2024. Characterization and functional analysis of Toxoplasma Golgi-associated proteins identified by proximity labeling. mBio, 15(11), pp e0238024.

Percie du Sert, N., Hurst, V., Ahluwalia, A., Alam, S., Avey, M. T., Baker, M., Browne, W. J., Clark, A., Cuthill, I. C., Dirnagl, U., Emerson, M., Garner, P., Holgate, S. T., Howells, D. W., Karp, N. A., Lazic, S. E., Lidster, K., MacCallum, C. J., Macleod, M., Pearl, E. J., Petersen, O. H., Rawle, F., Reynolds, P., Rooney, K., Sena, E. S., Silberberg, S. D., Steckler, T. & Würbel, H. 2020. The ARRIVE guidelines 2.0: Updated guidelines for reporting animal research. PLOS Biology, 18(7), pp e3000410.

Primo, V., A., Rezvani, Y., Farrell, A., Murphy, C., Q., Lou, J., Vajdi, A., Marth, G., T., Zarringhalam, K. & Gubbels, M.-J. 2021. The Extracellular Milieu of Toxoplasma’s Lytic Cycle Drives Lab Adaptation, Primarily by Transcriptional Reprogramming. mSystems, 6(6), pp e01196–21.

Ramakrishnan, S., Docampo, M. D., MacRae, J. I., Pujol, F. M., Brooks, C. F., van Dooren, G. G., Hiltunen, J. K., Kastaniotis, A. J., McConville, M. J. & Striepen, B. 2012. Apicoplast and endoplasmic reticulum cooperate in fatty acid biosynthesis in apicomplexan parasite *Toxoplasma gondii*. Journal of Biological Chemistry, 287(7), pp 4957–4971.

Ramakrishnan, S., Docampo, M. D., MacRae, J. I., Ralton, J. E., Rupasinghe, T., McConville, M. J. & Striepen, B. 2015. The intracellular parasite Toxoplasma gondii depends on the synthesis of long-chain and very long-chain unsaturated fatty acids not supplied by the host cell. Molecular microbiology, 97(1), pp 64–76.

Ren, B., Kong, P., Hedar, F., Brouwers, J. F. & Gupta, N. 2020. Phosphatidylinositol synthesis, its selective salvage, and inter-regulation of anionic phospholipids in Toxoplasma gondii. Communications Biology, 3(1), pp 750.

Rimple, P. A., Olafsson, E. B., Markus, B. M., Wang, F., Augusto, L., Lourido, S. & Carruthers, V. B. 2025. Metabolic adaptability and nutrient scavenging in *Toxoplasma gondii*: insights from ingestion pathway-deficient mutants. mSphere, 10(4), pp e01011–24.

Salman, D., Mahmoud, M. E., Pumidonming, W., Mairamkul, T., Oohashi, E. & Igarashi, M. 2021. Characterization of a spontaneous cyst-forming strain of Toxoplasma gondii isolated from Tokachi subprefecture in Japan. Parasitology International, 80(102199.

Sanchez, S. G., Bassot, E., Cerutti, A., Mai Nguyen, H., Aïda, A., Blanchard, N. & Besteiro, S. 2023. The apicoplast is important for the viability and persistence of Toxoplasma gondii bradyzoites. Proceedings of the National Academy of Sciences, 120(34), pp e2309043120.

Schindelin, J., Arganda-Carreras, I., Frise, E., Kaynig, V., Longair, M., Pietzsch, T., Preibisch, S., Rueden, C., Saalfeld, S., Schmid, B., Tinevez, J.-Y., White, D. J., Hartenstein, V., Eliceiri, K., Tomancak, P. & Cardona, A. 2012. Fiji: an open-source platform for biological-image analysis. Nature Methods, 9(7), pp 676–682.

Schmittgen, T. D. & Livak, K. J. 2008. Analyzing real-time PCR data by the comparative CT method. Nature Protocols, 3(6), pp 1101–1108.

Shears, M. J., MacRae, J. I., Mollard, V., Goodman, C. D., Sturm, A., Orchard, L. M., Llinás, M., McConville, M. J., Botté, C. Y. & McFadden, G. I. 2017. Characterization of the Plasmodium falciparum and P. berghei glycerol 3-phosphate acyltransferase involved in FASII fatty acid utilization in the malaria parasite apicoplast. Cellular Microbiology, 19(1), pp e12633.

Sheiner, L., Demerly, J. L., Poulsen, N., Beatty, W. L., Lucas, O., Behnke, M. S., White, M. W. & Striepen, B. 2011. A Systematic Screen to Discover and Analyze Apicoplast Proteins Identifies a Conserved and Essential Protein Import Factor. PLOS Pathogens, 7(12), pp e1002392.

Sheiner, L., Santos, J. M., Klages, N., Parussini, F., Jemmely, N., Friedrich, N., Ward, G. E. & Soldati-Favre, D. 2010. Toxoplasma gondii transmembrane microneme proteins and their modular design. Mol Microbiol, 77(4), pp 912–29.

Sheokand, P. K., Yamaryo-Botté, Y., Narwal, M., Arnold, C.-S., Thakur, V., Islam, M. M., Banday, M. M., Asad, M., Botté, C. Y. & Mohmmed, A. 2023. A Plasmodium falciparum lysophospholipase regulates host fatty acid flux via parasite lipid storage to enable controlled asexual schizogony. Cell Reports, 42(4), pp 112251.

Shunmugam, S., Arnold, C.-S., Dass, S., Katris, N. J. & Botté, C. Y. 2022. The flexibility of Apicomplexa parasites in lipid metabolism. PLOS Pathogens, 18(3), pp e1010313.

Smith, D., Kannan, G., Coppens, I., Wang, F., Nguyen, H. M., Cerutti, A., Olafsson, E. B., Rimple, P. A., Schultz, T. L., Mercado Soto, N. M., Di Cristina, M., Besteiro, S. & Carruthers, V. B. 2021. Toxoplasma TgATG9 is critical for autophagy and long-term persistence in tissue cysts. eLife, 10(e59384.

Srivastava, S., White Michael, W. & Sullivan William, J. 2020. Toxoplasma gondii AP2XII-2 Contributes to Proper Progression through S-Phase of the Cell Cycle. mSphere, 5(5), pp 10.1128/msphere.00542-20.

Suarez, C., Lentini, G., Ramaswamy, R., Maynadier, M., Aquilini, E., Berry-Sterkers, L., Cipriano, M., Chen, A. L., Bradley, P., Striepen, B., Boulanger, M. J. & Lebrun, M. 2019. A lipid-binding protein mediates rhoptry discharge and invasion in Plasmodium falciparum and Toxoplasma gondii parasites. Nature Communications, 10(1), pp 4041.

Tagoe, D. N. A., Ribeiro E Silva, A., Drozda, A. A., Coppens, I., Coleman, B. I. & Gubbels, M.-J. 2024. Toxoplasma FER1 is a versatile and dynamic mediator of differential microneme trafficking and microneme exocytosis. Scientific Reports, 14(1), pp 21819.

Tomita, T., Bzik, D. J., Ma, Y. F., Fox, B. A., Markillie, L. M., Taylor, R. C., Kim, K. & Weiss, L. M. 2013. The Toxoplasma gondii Cyst Wall Protein CST1 Is Critical for Cyst Wall Integrity and Promotes Bradyzoite Persistence. PLOS Pathogens, 9(12), pp e1003823.

Tymoshenko, S., Oppenheim, R. D., Agren, R., Nielsen, J., Soldati-Favre, D. & Hatzimanikatis, V. 2015. Metabolic Needs and Capabilities of Toxoplasma gondii through Combined Computational and Experimental Analysis. PLOS Computational Biology, 11(5), pp e1004261.

Venugopal, K., Werkmeister, E., Barois, N., Saliou, J.-M., Poncet, A., Huot, L., Sindikubwabo, F., Hakimi, M. A., Langsley, G., Lafont, F. & Marion, S. 2017. Dual role of the Toxoplasma gondii clathrin adaptor AP1 in the sorting of rhoptry and microneme proteins and in parasite division. PLOS Pathogens, 13(4), pp e1006331.

Waller, R. F., Keeling, P. J., Donald, R. G. K., Striepen, B., Handman, E., Lang-Unnasch, N., Cowman, A. F., Besra, G. S., Roos, D. S. & McFadden, G. I. 1998. Nuclear-encoded proteins target to the plastid in Toxoplasma gondii and Plasmodium falciparum. Proceedings of the National Academy of Sciences, 95(21), pp 12352–12357.

Walsh, D., Katris, N. J., Sheiner, L. & Botté, C. Y. 2022. Toxoplasma metabolic flexibility in different growth conditions. Trends in Parasitology.

Wang, Q., Du, X., Ma, K., Shi, P., Liu, W., Sun, J., Peng, M. & Huang, Z. 2018. A critical role for very long-chain fatty acid elongases in oleic acid-mediated Saccharomyces cerevisiae cytotoxicity. Microbiological Research, 207(1-7.

Wang, Q. & Sibley, L. D. 2020. Assays for Monitoring Toxoplasma gondii Infectivity in the Laboratory Mouse. In: Tonkin, C. J. (ed.) Toxoplasma gondii: Methods and Protocols. New York, NY: Springer US.

Wang, Y., Sangaré, L. O., Paredes-Santos, T. C., Hassan, M. A., Krishnamurthy, S., Furuta, A. M., Markus, B. M., Lourido, S. & Saeij, J. P. J. 2020. Genome-wide screens identify Toxoplasma gondii determinants of parasite fitness in IFNγ-activated murine macrophages. Nature Communications, 11(1), pp 5258.

Xia, N., Yang, J., Ye, S., Zhang, L., Zhou, Y., Zhao, J., David Sibley, L. & Shen, B. 2018. Functional analysis of Toxoplasma lactate dehydrogenases suggests critical roles of lactate fermentation for parasite growth in vivo. Cellular microbiology, 20(1), pp e12794.

Xia, N., Ye, S., Liang, X., Chen, P., Zhou, Y., Fang, R., Zhao, J., Gupta, N., Yang, S., Yuan, J., Shen, B., Vaidya, A. B. & Sibley, L. D. 2019. Pyruvate Homeostasis as a Determinant of Parasite Growth and Metabolic Plasticity in Toxoplasma gondii. MBio, 10(3), pp e00898–19.

Yang, Y., Yu, S.-M., Chen, K., Hide, G., Lun, Z.-R. & Lai, D.-H. 2020. Temperature is a key factor influencing the invasion and proliferation of Toxoplasma gondii in fish cells. Experimental Parasitology, 217(107966.

Zheng, T., Li, H., Han, N., Wang, S., Hackney Price, J., Wang, M. & Zhang, D. 2017. Functional Characterization of Two Elongases of Very Long-Chain Fatty Acid from Tenebrio molitor L. (Coleoptera: Tenebrionidae). Scientific Reports, 7(1), pp 10990.

